# An improved reference of the grapevine genome supports reasserting the origin of the PN40024 highly-homozygous genotype

**DOI:** 10.1101/2022.12.21.521434

**Authors:** Amandine Velt, Bianca Frommer, Sophie Blanc, Daniela Holtgräwe, Éric Duchêne, Vincent Dumas, Jérôme Grimplet, Philippe Hugueney, Marie Lahaye, Catherine Kim, José Tomás Matus, David Navarro-Payá, Luis Orduña, Marcela K. Tello-Ruiz, Nicola Vitulo, Doreen Ware, Camille Rustenholz

## Abstract

The genome sequence assembly of the diploid and highly homozygous *V. vinifera* genotype PN40024 serves as the reference for many grapevine studies. Despite several improvements of the PN40024 genome assembly, its current version PN12X.v2 is quite fragmented and only represents the haploid state of the genome with mixed haplotypes. In fact, despite the PN40024 genome is nearly homozygous, it still contains various heterozygous regions. Taking the opportunity of the improvements that long-read sequencing technologies offer to fully discriminate haplotype sequences and considering that several *Vitis* sp. genomes have recently been assembled with these approaches, an improved version of the reference, called PN40024.v4, was generated.

Through incorporating long genomic sequencing reads to the assembly, the continuity of the 12X.v2 scaffolds was highly increased. The number of scaffolds decreased from 2,059 to 640 and the number of N bases was reduced by 88%. Additionally, the full alternative haplotype sequence was built for the first time, the chromosome anchoring was improved and the amount of unplaced scaffolds were reduced by half. To obtain a high-quality gene annotation that outperforms previous versions, a liftover approach was complemented with an optimized annotation workflow for *Vitis*. Integration of the gene reference catalogue and its manual curation have also assisted in improving the annotation, while defining the most reliable estimation to date of 35,230 genes. Finally, we demonstrate that PN40024 resulted from selfings of cv. ‘Helfensteiner’ (cross of cv. ‘Pinot noir’ and ‘Schiava grossa’) instead of a single ‘Pinot noir’. These advances will help maintaining the PN40024 genome as a gold-standard reference also contributing in the eventual elaboration of the grapevine pangenome.

## Introduction

Cultivated grapevine (*Vitis vinifera* ssp. *vinifera*) was the fourth plant whose genome was sequenced and assembled ^1^. Because of the grapevine’s high level of heterozygosity (one Single Nucleotide Polymorphism (SNP) per 100 bp and one Indel per 450 bp, ^2^), the genotype selected for sequencing was PN40024, whose ~475 Mb genome ^3^ is near homozygous (estimated at ~93%). PN40024 was indeed generated through nine rounds of selfing and supposedly originated from ‘Pinot noir’, hence its identification as ‘PN’. This unique genome characteristic allowed a high-quality whole-genome shotgun assembly based on 8X coverage Sanger reads ^1^. In 2009, a 4X coverage was added, which improved the overall coverage of the genome (from 68.9% for the 8X version to 91.2% for the 12X.v0) (http://urgi.versailles.inra.fr/Species/Vitis/Data-Sequences/Genome-sequences; FN597015-FN597047 at EMBL, release 102; Supplementary File 1 Fig. S1). In 2017, a third assembly version, named 12X.v2, was published as the result of a large anchoring effort using six dense parental genetic maps ^4^ Despite these advances, no additional sequencing efforts have been made and although it’s very high quality, the 12X.v0 Sanger contigs are numerous (14,642), the 12X.v2 scaffolds are composed of large N gaps (3.1% of the cumulative scaffold size) and the 19 pseudomolecules are quite fragmented (19.3 scaffolds on average per pseudomolecule).

In the last years the advent of third generation sequencing technologies, especially those from the Pacific Biosciences (PacBio) platform, have allowed the assembly of grapevine diploid genomes with a higher level of contiguity compared to the 12X.v2 version of the PN40024 genome (for example, cv. ‘Cabernet Sauvignon’ genome assembly ^5^).

Along with the versions of each genome assembly, several versions of gene annotations were made available (Supplementary File 1 Fig. S1). The first version of the grapevine genome assembly, 8X, was published along with the prediction of 30,434 gene models based on the GAZE software ^1,6^. For the 12X.v0, three different versions of gene predictions were made available: the v0 version (26,346 gene models), based on the GAZE software ^6^, the CRIBIv1 version (29,971 gene models), based on the JIGSAW software ^7^, and the CRIBIv2 version (31,845 gene models), with an effort made on the discovery of splicing variants ^8^. For the 12X.v2, the International Grapevine Genome Program (IGGP) led the initiative of merging annotations from NCBI Refseq, CRIBIv1 and VCost, which was based on the Eugene software ^9^ and was generated in the frame of the COST Action FA1106. This version, called VCost.v3, resulted in an exhaustive view of the PN40024 grapevine gene content with its 42,413 gene models ^4^. However, after several years as the reference annotation by the grapevine scientific community, it appeared that the great increase in number of gene models for VCost.v3 compared to all the previous annotation versions was caused by many small and fragmented predictions that were probably erroneous.

By combining the top-quality Sanger contigs from the 12X version and long reads generated here by Single-Molecule Real-Time (SMRT) sequencing (PacBio), we provide an improved version of the PN40024 genome sequence assembly, referred to as PN40024.v4. Along with this new assembly, we also provide a new version of the gene annotation, PN40024.v4.2, based on a newly-developed annotation workflow, RNA-Seq datasets and an exhaustive manual curation of a set of catalogued genes of functional interest to the community. Finally, we demonstrate that PN40024 originates from selfings of the ‘Helfensteiner’ cultivar instead of ‘Pinot noir’.

## Material and Methods

### Plant material, DNA extractions and sequencing

DNA extractions of young leaves of cv. ‘Pinot noir’ clone 162 (ID code FRA038-193.Col.162), cv. ‘Schiava grossa’ (synonymous ‘Trollinger’, ID code FRA038-2525.Col.1) and cv. ‘Helfensteiner’ (ID code FRA038-2744.Col.1) were performed as described by Merdinoglu *et al*. ^10^. Illumina DNA PCR-Free Prep kit was used to prepare the resequencing libraries according to provider’s procedure. Paired-end Illumina HiSeq 4000 sequencing at about 15x coverage was performed for ‘Pinot noir’ and ‘Schiava grossa’, respectively. Paired-end Illumina NovaSeq 6000 sequencing at about 15x coverage was performed for ‘Helfensteiner’.

One gram of young leaves (1cm^2^) of PN40024 (ID code FRA038-40024.Col.1) was collected and DNA was extracted using QIAGEN Genomic-tips 100/G kit. SMRT sequencing on a Sequel I machine (3 SMRTCells; PacBio) and dedicated library preparation were performed according to provider’s procedure.

Genotyping-by-sequencing (GBS) was performed on the population ‘Riesling’ x ‘Gewurztraminer’ (exhaustively described by ^11^ using the procedure described by Girollet *et al*. ^12^.

All data generated in the frame of this study have been submitted under the ENA Study Accession PRJEB45423.

### Genome assembly

Raw SMRT reads (ERR7997743) were self-corrected using CANU (v.1.6) ^13^, followed by a correction with PN40024 Illumina reads (SRR8835144) using LORDEC (v.0.5.3) ^14^ The corrected reads were mapped on PN12X scaffolds (https://urgi.versailles.inra.fr/download/vitis/VV_12X_embl_102_Scaffolds.fsa.zip) using minimap2 (v2.17-r954-dirty) ^15^. A total of 163,446 reads (15%) were aligned on less than 80% of their length and/or with less than 80% identity and were thus considered as missing from PN12X scaffolds. These unmapped reads were assembled using Flye (v2.4-gc9db046) ^16^. We aligned these new contigs on the Uniprot *Arabidopsis* database (release 2019_01) using blastx ^17^. Contigs longer than 5 kb and having hit(s) with *Arabidopsis* proteins with >60% identity and >60% length coverage, were selected for the next step. The fasta files of these new contigs and the PN12X scaffolds were concatenated to generate the new assembly. Firstly, the repeats were masked using Red (v05/22/2015) ^18^. Then, Haplomerger2 (v20180603) ^19^ was used following three steps according to developer’s procedure: i) break the misjoins and output the new diploid assembly; ii) separate/merge two haplotypes and output haploid assemblies (REF and ALT); and iii) remove tandem errors from haploid assemblies. Some scaffolds / contigs were deleted by Haplomerger2 during the assembly process but sequences longer than 10 kb were retrieved and added to the REF scaffolds. The two haploid assemblies (REF/ALT) were then scaffolded with the OPERA-LG tool (v2.0.6) ^20^, which uses both, corrected SMRT reads and Illumina reads. A first gap-filling step (two rounds) was carried out with Illumina reads using GapCloser (v1.12) ^21^ and a second gapfilling step (three rounds) was carried out with corrected SMRT reads using LR_Gapcloser (v1.0) ^22^. A final polishing step was performed with the Illumina reads using PILON (v1.23-1-g41e0b8e) ^23^ (Fig. 1A and 1B).

**Fig. 1:**
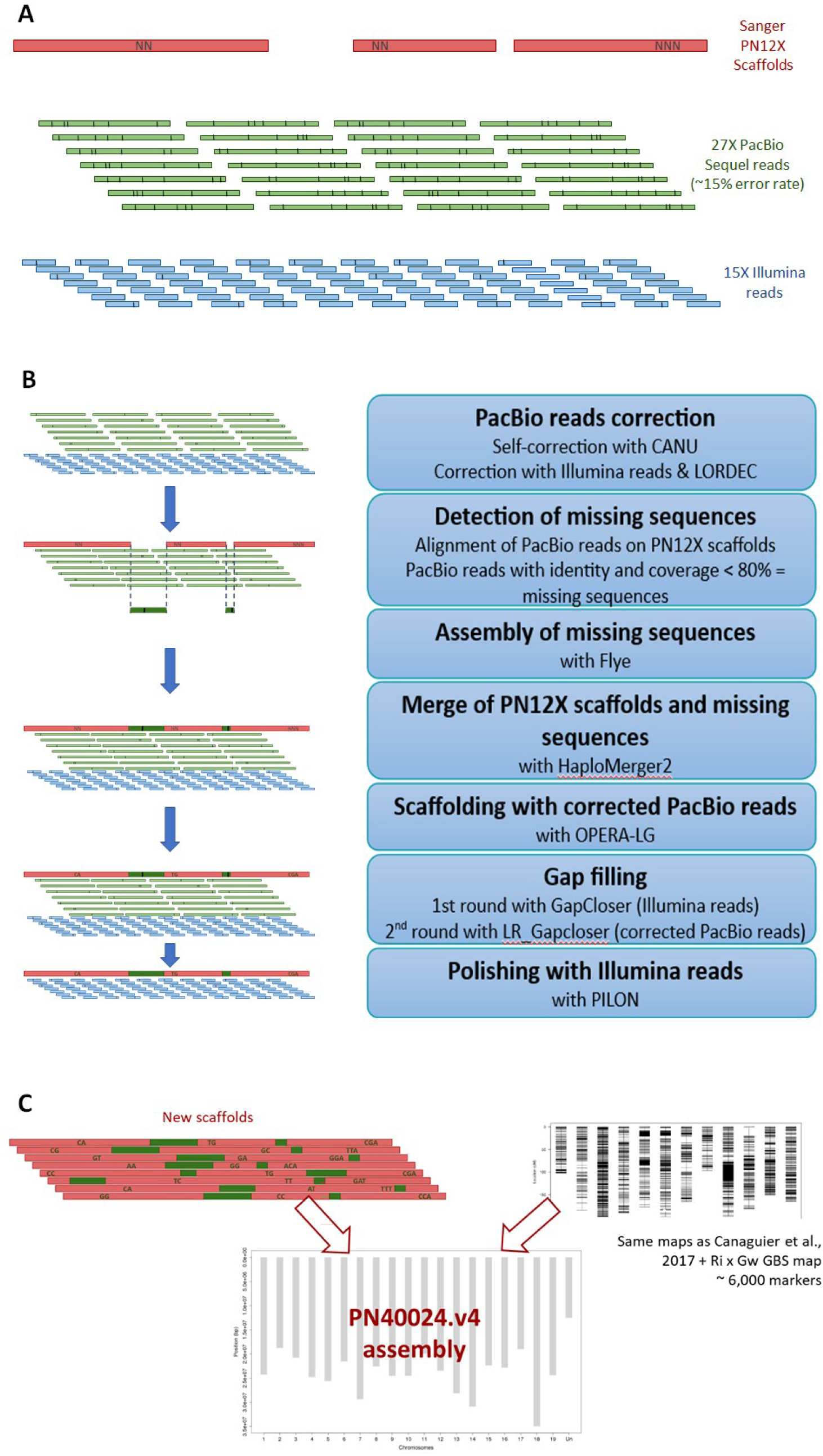
Assembly process for the PN40024.v4 genome sequence assembly. **A**) Initial datasets: Sanger-based scaffolds of PN12X.v2 (red) with unknown bases (‘N’s’), genomic SMRT reads (green) and genomic short reads (blue). Erroneous bases are represented by a black line. **B**) Scaffold assembly steps. Same color code as A). Dark green regions represent newly incorporated SMRT sequencing regions. **C**) Pseudomolecule construction using the new scaffolds and genetic maps. Sequence region of 12X.v2 are colored red and newly incorporated SMRT sequencing regions are colored dark green.

The anchoring of the new haploid scaffolds was performed using the six genetic maps used for the same purpose by ^4^ and two new genetic maps from cv. ‘Riesling’ and cv. ‘Gewurztraminer’ derived from GBS. To transfer the markers from ^4^ from PN12X.v2 to the scaffolds of PN40024.v4, BLAST (v2.2.28) ^17^ or ipcress (ipcress from exonerate v2.2.0) ^24^ was used to align the markers and find the position of each on the scaffolds of PN40024.v4 REF and ALT. A total of 2,333 markers for REF and 2,326 markers for ALT were used from these six maps to anchor the scaffolds. For the two new genetic maps from ‘Riesling’ and ‘Gewurztraminer’, 5,884 (‘Riesling’) and 5,840 (‘Gewurztraminer’) SNP markers were available for REF and 5,866 (‘Riesling’) and 5,832 (‘Gewurztraminer’) for ALT. The SNP markers were derived from GBS data (ERR8657388 to ERR8657647) and were analyzed with Fast-GBS ^25^ with modifications to allow paired-end read analysis (https://forgemia.inra.fr/sophie.blanc/gbs). The two genetic maps were built using R ASMap package with the “kosambi” parameter ^26^. A first run of Allmaps (v0.9.13) ^27^ was performed with the “merge” command to merge all genetic maps and then “split”, “gaps”, “refine” and “build” commands to create breakpoints (58 for REF scaffolds and 47 for ALT scaffolds), with default parameters. Subsequently, all maps were recreated for new scaffolds and then orientation and anchoring of new haploid scaffolds on the 19 pseudochromosomes was performed using Allmaps with the “merge” command to merge all maps and “path” command to anchor, with default parameters (Fig. 1C).

### Quality assessment of the PN40024.v4 genome sequence assembly

A quality analysis of the genome assembly was done with Merqury v1.3 ^28^. Since PN40024 is a ‘Helfensteiner’ selfing (demonstrate below) and since ‘Helfensteiner’ originated from a cross between ‘Pinot noir precoce’ and ‘Schiava grossa’, ‘Schiava grossa’ was used as the maternal parent. The run was carried out on the scaffolds using genomic paired-end short reads of PN40024 as the child data (SRR8835144), short reads of ‘Pinot noir’ as the paternal data (ERR8014965) and short reads of cv. ‘Schiava grossa’ as the maternal data (ERR8014964). A k-mer database was built for the three read datasets with k=19, the Merqury hap-mer databases were computed and the PN40024.v4 genome assembly was evaluated using ‘num_switch 100’ and ‘short_range 20,000’. For comparison reasons, the Merqury quality analysis was carried out on PN12X.v2 using the same k-mer databases.

The “*Flowering locus T*” (*FT*) and the “*Adenine phosphoribosyltransferase 3*” (*APRT3*) genes are absent and truncated in PN12X.v2, respectively. To check whether these genes could be retrieved in the new genome assembly, cDNA sequences of *FT* (NM_001280978.1) and *APRT3* (GSVIVT00007310001, PN8X version) were used to perform blastn ^17^ against PN8X, PN12X, PN12X.v2 and PN40024.v4 genome assemblies. High Scoring Pairs (HSPs) were then accumulated for each analysis and the mean percentage identity, query overlap, hit query start and end were calculated.

PN40024 (SRR8835144), and the cultivars ‘Silvaner Gruen’ (SRR5891620), ‘Cabernet Franc’ (SRR5891774), ‘Cabernet Sauvignon’ (SRR5891776), ‘Chardonnay’ (SRR5891778), ‘Muscat Hamburg’ (SRR5891787), ‘Semillon’ (SRR5891866), ‘Pinot Noir’ (SRR5891886), ‘Merlot’ (SRR5891890), ‘Sauvignon Blanc’ (SRR5891893), ‘Muscat of Alexandria’ (SRR5891985) and ‘Riesling’ (SRR5891989) genomic paired-end resequencing datasets were aligned against PN40024.v4 REF, PN12X.v2 and ‘Cabernet Sauvignon’ haplotype 1 ^5^ pseudomolecule assemblies (without chrUn or unplaced contigs / scaffolds) using bwa-mem2 (v2.0) ^29^ “mem” command with default parameters. Samtools’ (v1.9) ^30^ “flagstat” command was used with default parameters to compute alignment statistics.

PN12X scaffolds were mapped against PN40024.v4 REF pseudomolecules using NUCmer (MUMmer v3.1) ^31^ with “-maxmatch -l 100 -c 500” parameters. The output file was filtered using MUMmer show-coords command with “-l -g -I 99.5” parameters. The resulting file was formatted into BED format and merged with the bed file corresponding to N gap regions in the PN40024.v4 assembly. Pseudomolecule regions over 100 bp that did not correspond to either PN12X scaffolds or N gap regions were identified as ‘newly assembled’ PacBio long read-based regions.

The identification of variants between PN40024 paired-end Illumina resequencing (SRR8835144) and PN40024.v4 REF and ALT pseudomolecules was performed as described in section “Origin of PN40024”. The homozygous calls “1/1” were considered as assembly errors. The densities of the heterozygous calls “0/1” along the REF and ALT pseudomolecules were used to define seven heterozygous regions of the PN40024 genome.

### Origin of PN40024

PN40024 (SRR8835144), ‘Pinot noir’ (ERR8014965), ‘Schiava grossa’ (ERR8014964), ‘Helfensteiner’ (ERR8014963) and ‘Araklinos’ (SRR8835172) paired-end resequencing datasets were all analyzed using the same pipeline. Datasets were aligned against PN40024.v4 REF assembly using bwa-mem2 (v2.0) ^29^ “mem” command with default parameters. Samtools (v1.9) ^30^ “view” and “sort” commands with default parameters were used to convert and sort the output BAM files. GATK (v4.1.4.0) ^32^ “MarkDuplicatesSpark”, “HaplotypeCaller” and “GenotypeGVCFs” commands with default parameters were used to generate variant files in VCF format. The GATK “VariantFiltration” command was used to filter out variants meeting at least one of the following criteria: QD < 8.0, QUAL < 100.0, FS > 60.0, SOR > 3.0, DP < 3, DP > 30, AD < 2. The final variant files were obtained using GATK “SelectVariants” command with “--exclude-filtered --exclude-non-variants” parameters. The homozygous SNP calls “1/1” were selected for each analyzed genotype. All SNPs corresponding to a homozygous call in PN40024 genotypes were excluded from the analysis as they represent assembly errors. The remaining homozygous SNPs were used to draw density plots on the PN40024.v4 pseudomolecules. The regions that are rich in homozygous SNPs for a given genotype correspond to regions for which this genotype does not share a haplotype with PN40024.

The haplotypic blocks were defined after segmentation of homozygous SNP densities along the chromosomes using the R package changepoint (v2.2.2) ^33^ with command “cpt.mean” and the parameters method=“PELT” and penalty=“AIC”. Some manual curation of the segments was performed to join directly adjacent segments of the same origin (‘Pinot noir’ or ‘Schiava grossa’). The size of the segments was used to calculate the proportion of ‘Pinot noir’, ‘Schiava grossa’ and common haplotypes.

### Gene prediction

Before performing gene prediction, the PN40024.v4 genome assembly was repeat masked with RepeatMasker v4.1.2 ^34^ using crossmatch as search engine. Predictions with a SW-Score <1,000 were filtered out and predictions with a Smith-Waterman (SW)-Score between 1,000 and 2,000 were only kept if the reported percentage of substitutions were <20%. The PN40024.v4 genome assembly was softmasked with BEDTool (v2.26.0) ^35^.

To annotate the PN40024.v4 genome assembly, publicly available *V. vinifera* stranded (Supplementary File 2 Table S1) and unstranded (Supplementary File 2 Table S2) paired-end RNA-Seq datasets of different tissues and treatments were collected. RNA-Seq data were trimmed with Trimmomatic (v0.39) ^36^. The annotation pipeline was first tested on the PN40024 12X.v0 genome assembly using VCost.v3 gene annotation as quality reference. The gene predictors SNAP ^37^ and BRAKER2 ^38–40^ were trained and tested on the softmasked 12X.v0 genome assembly. The RNA-Seq data were mapped on 12X.v0 and on PN40024.v4 REF and ALT sequences with GMAP/GSNAP v2020-09-12 setting “-B 5 --novelsplicing 1” ^41^. Primary mappings were extracted with SAMTools v1.9 ^30^. Based on the primary mappings, stranded and unstranded reference-guided transcriptome assemblies were computed with PsiCLASS v1.0.1 using default parameters ^42^.

Additionally, *A. thaliana* protein sequences (UniProt/SwissProt release 2020_02), eudicotyledone protein sequences (UniProt/SwissProt release 2020_02, OrthoDB10 v1), Viridiplantae and Vitales sequences (UniProt/SwissProt release 2020_02) were aligned on 12X.v0 and on PN40024.v4 REF and ALT with pBLAT v1.9 ^43^, a parallel implementation of the original blat algorithm ^44^. The genome regions on which the protein data mapped were extracted and the protein sequences were aligned to these regions with exonerate v2.4.0 ^24^ Only the proteins that aligned on the reference genome with an identity of 25%, a similarity of 50% and with a sequence alignment coverage of at least 80%, were retained and included in the gene prediction.

The gene predictor GlimmerHMM v3.0.4 ^45^ was trained on 12X.v0 and on PN40024.v4 REF and ALT using 7,500 (12X.v0) and 15,000 (PN40024.v4) random PsiCLASS transcripts of the 12X.v0 or PN40024.v4 REF or ALT stranded transcriptome assembly, respectively. The training was followed by gene prediction with GlimmerHMM with default settings.

Moreover, the gene predictor SNAP v2006-07-28 was trained on the 12X.v0 genome assembly. For this, the 12X.v0 genome assembly, the stranded transcriptome assembly, the Viridiplantae protein sequences and the eudicotyledone protein sequences were given to MAKER2 v3.01.03 ^46,47^ and initial data alignment with BLAST (ncbi-blast-2.10.1+) ^17,48^ and exonerate was performed followed by MAKER2 *ab initio* gene prediction. MAKER2 was run with “max_dna_len=300000” and “split_hit=20000”. A SNAP hmm file was generated with the MAKER2 gff file and a second MAKER2 run was performed with enabled SNAP gene prediction and the SNAP hmm file as input. Hmm file generation and SNAP gene prediction with MAKER2 and the new hmm file was repeated. The hmm file generated with the 12X.v0 assembly was used to run SNAP gene prediction on the PN40024.v4 REF and ALT genome sequences.

An AUGUSTUS species model was computed with BRAKER2 v2.1.5-master_20200915 and the 12X.v0 genome assembly. BRAKER2 was run with enabled softmasking and in *etpmode* calling GeneMark-ETP+ v4.61 ^49–51^ for initial gene prediction followed by AUGUSTUS training and gene prediction (AUGUSTUS version master_v3.3.3_20200914) ^52,53^. With BRAKER2, the programs DIAMOND v0.9.24.125 ^54^, SAMtools v1.9-180-gf9e1caf ^30^, SPALN version 2.3.3f ^55,56^, ProtHint version 2.5.0 and BamTools v2.5.1 ^57^ were called. The stranded RNA-Seq primary mappings, the eudicotyledon protein sequences (OrthoDB10 v1) and the Viridiplantae protein sequences were used as input. The gene prediction on PN40024.v4 REF and ALT was performed with BRAKER2 v2.1.5-master_20210218, the generated AUGUSTUS species model and AUGUSTUS version master_v3.4.0_20210218. Again, the stranded RNA-Seq mappings and the same protein sequences were used as input. The BRAKER2 parameter settings were left the same as above.

The last *ab initio* gene prediction was done on the PN40024.v4 genome assembly with GeneID v1.4.5-master-20200916 and the publicly available *V. vinifera* parameter set using default settings. To add the VCost.v3 gene annotation to the set of predictions, an annotation liftover was performed with liftoff v1.5.1 ^58^ with default parameters onto the PN40024.v4 genome assembly.

To combine the predictions and evidence data into an overall gene model set, the GlimmerHMM, SNAP, BRAKER2 and GeneID *ab initio* gene prediction as well as the lifted VCost.v3 annotation, the stranded and unstranded transcriptome assemblies, the GFF file with the aligned protein data, the repeat annotation GFF file and the PN40024.v4 genome assembly was given to EvidenceModeler v1.1.1 ^59^. The used weights are listed in Supplementary File 2 Table S3.

Subsequently, the raw gene models were quality filtered. Gene models only supported by *ab initio* predictors were kept if at least two gene prediction programs predicted them, if the start and stop codon was present and if the gene length was equal or larger than 300 bp. However, *ab initio* supported gene models not matching these constraints were kept if they had a database hit with the UniProt/SwissProt or NCBI non-redundant database. To obtain that, a blastp search of the protein sequences against the two databases was run, allowing hits with an e-value <1e^-6^. Of the gene models only supported by evidence data or by VCost.v3 lifted annotation, those gene models with missing start and stop and a gene length <300 bp were discarded.

The gene models generated by EvidenceModeler were finally processed by PASA (v2.4.1, default parameters) using the stranded transcriptome assembly as a reference to add UTR regions and to calculate alternatively spliced models. Genes with overlapping UTRs were shortened. tRNAs were predicted with tRNAscan-SE-2.0 ^60^ on the PN40024.v4 genome assembly.

To retain gene naming of VCost.v3 gene models, a reciprocal best blast hit (RBH) search between protein sequences of PN40024.v4.1 gene models and protein sequences of VCost.v3 gene models was carried out. For the RBH search, only the longest protein sequence per gene was used, the e-value was set to 1e^-4^ and the query coverage and identity was set to 70%. Moreover, only RBHs with genes on the same pseudochromosome and showing collinearity with other genes were considered valid. Thus, genes with a valid RBH were named according to the VCost.v3 gene, novel genes received the prefix ‘04’ at the start of the gene number and genes predicted for alternative heterozygous sequence regions received the suffix ‘_alt’ (Supplementary File 2 Table S4).

The PN40024.v4.1 gene models were functionally annotated with Blast2GO (v1.5.1) ^61,62^. For this, protein domains of the PN40024.v4.1 proteins were identified with InterProScan (v5.52-86.0) ^63^ with options “--goterms --pathways-dp” using the databases/tools CDD-3.18 ^64^, Coils-2.2.1 ^65^, Gene3D-4.3.0 ^66^, Hamap-2020_05 ^67^, MobiDBLite-2.0 ^68^, PANTHER-15.0 ^69^, Pfam-33.1 ^70^, PIRSF-3.10 ^71^, PIRSR-2021_02, PRINTS-42.0 ^72^, ProSitePatterns-2021_01, ProSiteProfiles-2021_01 ^73^, SFLD-4 ^74^, SMART-7.1 ^75^, SUPERFAMILY-1.75 ^76,77^, TIGRFAM-15.0 ^78^. PN40024.v4.1 protein sequences were aligned with diamond “blastp” (v2.0.11) ^54^ to the NCBI nr database (nr.07_07_2021.fasta) with options “--sensitive --top 5 -e 1e-5 -f 5”. InterProScan and diamond results were used as input for Blast2GO.

### Quality assessment of the PN40024.v4.1 gene annotation

To estimate completeness of the PN40024.v4.1 gene model set, plant core genes were predicted with BUSCO v5.1.2 ^79,80^ using database eudicots_odb10.

Samples previously analyzed by ^81^ were used to perform differential gene expression analysis by using either PN12X.v2 assembly with VCost.v3 annotations or PN40024.v4 assembly with PN40024.v4.1 annotations. We analyzed cv. ‘Sangiovese’ (SRR1631822; SRR1631823; SRR1631824), cv. ‘Barbera’ (SRR1631834; SRR1631835; SRR1631836) and cv. ‘Refosco’ samples (SRR1631858; SRR1631859; SRR1631860) for the stage “Berries beginning to touch” (~EL35 according to Eichhorn and Lorenz phenological scale ^82^). The RNA-Seq data were downloaded from the SRA with SRA Toolkit (v2.10.8) (SRA Toolkit Development Team; https://trace.ncbi.nlm.nih.gov/Traces/sra/sra.cgi?view=software) and analyzed with an *in-house* pipeline using FASTQC (v0.11.5) ^83^, STAR (v2.5.3a) ^84^, Samtools (v1.4.1) ^30^, Bamtools (v2.4.0) ^57^, featureCounts (v1.5.3) ^85^ and SARTools (v1.7.3) ^86^.

### Manual gene model curation

For manual gene model curation, an Apollo Webserver v2.6.4 (https://github.com/GMOD/Apollo/blob/master/README.md) ^87^ was set up for the PN40024.v4 genome assembly and provided with different data tracks such as the PN40024.v4.1 and previous gene annotations, RNA-Seq mappings and exonerate protein mappings (see section Material and Methods - Gene prediction). By these means, gene models were manually inspected and curated if needed or also new genes were added following dedicated guidelines offered to the community (https://integrape.eu/resources/data-management/). Using Apollo, the plant core genes classified as fragmented and missing by BUSCO were manually curated and adapted if necessary. In the frame of this study, we also begun to manual curation of genes present in the grape reference catalogue (^88^; https://grapedia.org/genes/). A home-made python script was used to generate the PN40024.v4.2 version of gene annotations including manually curated ones (https://gitlab.com/MSVteam/pn40024-visualization-tools/-/tree/master/update_gff3_script).

## Results and Discussion

### Improved metrics for the genome assembly of PN40024

A hybrid strategy was developed to assemble the genome of PN40024 genotype using 27X of SMRT long reads along with the PN12X scaffolds and 15X PN40024 Illumina paired-end resequencing data (Fig. 1). This new assembly was named PN40024.v4. Six hundred and forty scaffolds were produced with a N50 size of 6.5 Mb for a cumulative size of 474.5 Mb for the PN40024.v4 REF haplotype (Table 1). Compared to the former PN12X.v2, the number of scaffolds was reduced by a factor of three and the N50 was doubled. Moreover, the number of unknown bases, marked as N in the new scaffold sequences, represents 1.8 Mb and 0.4% of the assembly size versus 15.0 Mb and 3.1% for PN12X.v2 scaffolds. Thus, PN40024.v4 REF is more contiguous and has more informative sequences than PN12X.v2. Also, the PN40024.v4 assembly size is closer to the grapevine genome size of 475 Mb, estimated using flow cytometry by Lodhi and colleagues ^3^. Phasing efforts on the partially heterozygous genotype resulted in the reconstruction of the second PN40024 haplotype (PN40024.v4 ALT) with 485 scaffolds and a total genome assembly size of ~463 Mb (Supplementary File 2 Table S5). Thus, the PN40024.v4 genome assembly now represents both haplotypes of the diploid PN40024 genome.

**Table 1:** Assembly statistics of the PN40024.v4 REF and PN12X.v2 genome assembly. The table lists statistics of the PN40024.v4 REF and PN12X.v2 scaffolded and chromosome-anchored genome assemblies. ‘N’ denotes the number (No.) of unknown bases.

| SCAFFOLDS | No. Scaf. | Min. size [bp] | Avg. size [kb] | Median size [kb] | L5 0 | N50 [Mb] | Max. size [Mb] | Sum [Mb] | No. Ns | GC [%] |
| --- | --- | --- | --- | --- | --- | --- | --- | --- | --- | --- |
| **PN12X.v2** | 2,059 | 2,001 | 236 | 5 | 41 | 3.43 | 13.10 | 485.19 | 14,976,411 | 33.5 |
| **PN40024.v4** | 640 | 542 | 742 | 20 | 25 | 6.50 | 15.23 | 474.61 | 1,755,062 | 34.4 |
| **Anchored PN12X.v2** | 367 | 2,010 | 1,250 | 277 | 37 | 3.57 | 13.10 | 465.64 | 11,921,253 | 33.6 |
| **Anchored PN40024.v4** | 165 | 1,085 | 2,801 | 1,506 | 24 | 6.57 | 15.23 | 462.14 | 1,631,047 | 34.4 |

There are 7,640 newly assembled PacBio long read-based regions that were identified as missing from PN12X.v2 scaffolds. Their cumulative size is 24.1 Mb, *i.e*. 5.1% of the total PN40024.v4 genome assembly size (average = 3,152 bp; median = 558 bp; max = 32,650 bp).

A total of 2,333 markers were used from the six Canaguier’s maps ^4^, in addition to 5,884 and 5,840 SNP markers from cv. ‘Riesling’ and cv. ‘Gewurztraminer’ GBS maps, respectively, to anchor these scaffolds. We were able to anchor 165 PN40024.v4 REF scaffolds to the 19 pseudochromosomes, for a cumulative size of ~462 Mb (97.4%) (Table 1). The 19 PN40024.v4 REF pseudomolecules are composed of 8.7 scaffolds on average (min = 3; max = 26; median = 6) whereas 19.3 scaffolds on average composed the PN12X.v2 pseudomolecules (min = 5; max = 82; median = 13). The remaining unplaced scaffolds were ordered according to their size to generate “chrUn” sequence representing 12.5 Mb (−47% compared to PN12X.v2 unplaced scaffolds). Thus, PN40024.v4 anchoring was improved as the pseudomolecules are less fragmented and as the size of chrUn has almost been halved.

At the chromosome scale, 10 pseudomolecules became shorter in PN40024.v4 compared to PN12X.v2 (average loss = ~448 kb; median = ~255 kb; min = 2,961 bp; max = 1,133,439 bp). Chromosome 6 showed the biggest reduction as the location of a large fragment has been correctly assigned to chromosome 9 (Supplementary File 1 Fig. S2). Nine pseudomolecules became larger (average gain = ~869 kb; median = ~582 kb; min = 15,118 bp; max = 2,045,414 bp), notably chromosome 9, 7 and 15, which gained 1.5 Mb, 1.9 Mb and 2.0 Mb, respectively.

By aligning PN40024 Illumina paired-end reads against PN40024.v4 genome assembly, we identified 101,778 heterozygous variations. Using their density along the chromosomes, we were able to identify seven well defined heterozygous regions in PN40024.v4 genome assembly as it was the first time that a software dedicated to diploid assembly (Haplomerger2) was used to assemble the PN40024 genome. These regions were located on chromosomes 2, 3, 4, 7, 10, 11 and 15 with the two largest regions being on chromosome 7 and 10 (11.4 Mb and 5.5 Mb, respectively) (Fig. 3). Their overall cumulative size of 20.6 Mb represents 4.3% of PN40024.v4, which is less than the residual heterozygosity size of 7%, estimated by Jaillon and colleagues based on genetic markers ^1^. Using the same procedure, we identified six heterozygous regions in PN12X.v2 assembly on the same chromosomes as PN40024.v4 except for the one on chromosome 15. Their overall cumulative size of 16.6 Mb represents 3.4% of PN12X.v2 and 4 Mb less than the heterozygous regions anchored on the PN40024.v4 chromosomes. These sequences were badly resolved and mostly located in the unanchored fraction of PN12X.v2 assembly (Supplementary File 1 Fig. S2). Thus, we conclude that PN40024.v4 is a better diploid assembly compared to PN12X.v2.

**Fig. 2:**
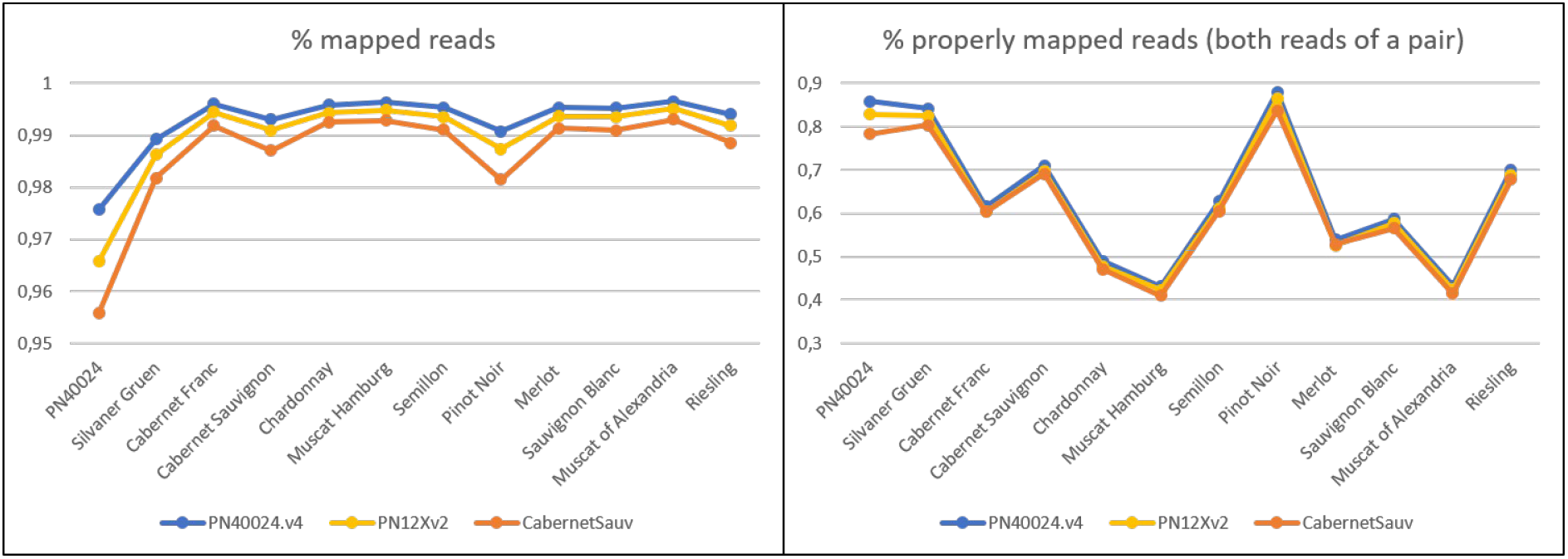
Percentage of mapped genomic reads and percentage of properly paired genomic reads between PN40024.v4, PN12X.v2 and cv. ‘Cabernet Sauvignon’ (Massonet et al., 2020) for 11 paired-end resequencing datasets of *V. vinifera* cultivars. The x-axis denotes the source (cultivar) of the genomic reads and the y-axis the percentage [%] of mapped reads. Note that the PN40024 dataset was obtained with Illumina Genome Analyzer IIx sequencing and all other samples with Illumina HiSeq 4000. The PN40024 dataset is therefore of lower quality than the others.

**Fig. 3:**
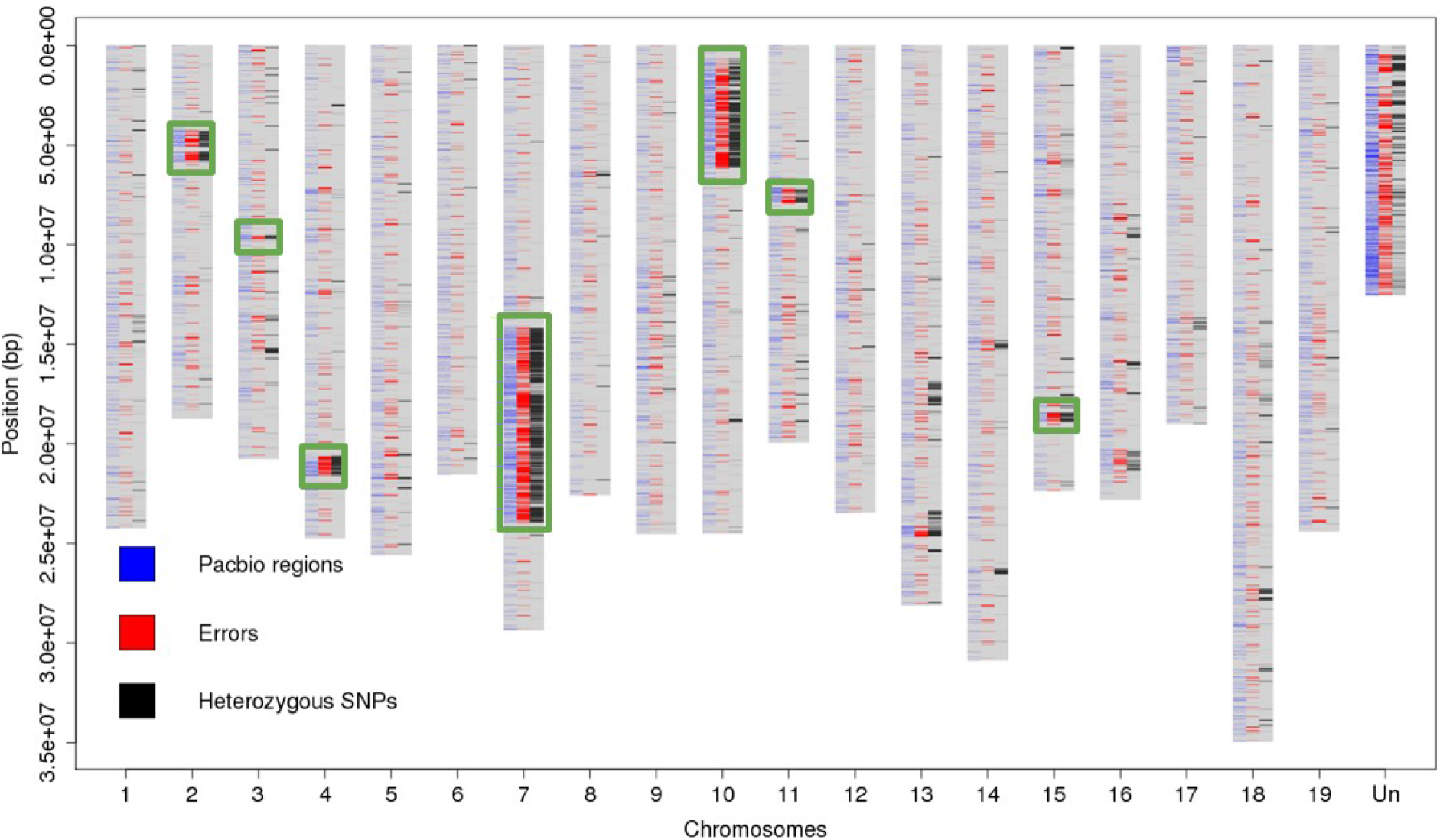
Location of regions assembled using long reads, density of errors and of heterozygous SNPs in the PN40024.v4 genome sequence assembly. The x-axis shows the 19 main pseudochromosomes and the artificial chrUn (‘Un’). The y-axis shows the base position in [bp]. ‘Pacbio regions’ refers to sequences derived from genomic SMRT reads. The seven heterozygous regions are squared in green.

### Quality of the PN40024.v4 genome assembly

The BUSCO analysis performed on the PN40024.v4 genome assembly confirmed that the gene space was more complete with 98.1% of the 2,326 total searched Eudicots BUSCO genes being complete, compared to PN12X.v2 with 97.6% (Fig. 6). The *FT* (*Flowering locus T*) gene is conserved among all flowering plants as it promotes transition from vegetative growth to flowering. However, its sequence could only be found on an unanchored scaffold in the PN8X version and was totally missing in PN12X.v0 and PN12X.v2. It is now present on chromosome 7 of the PN40024.v4 assembly and also on its allelic region, chromosome 7_ALT sequence. Similarly, the *APRT3* gene, located in the sex determination locus of grapevine, was present on chromosome 2 in the PN8X version and was truncated in PN12X.v0 and PN12X.v2. It is now be fully retrieved on chromosome 2 of PN40024.v4 assembly and on its allelic region, chromosome 2_ALT sequence. These two examples, along with the BUSCO analysis, show that the PN40024.v4 assembly is more complete, especially in the residual heterozygous regions that are now more accurately exposed.

**Fig. 4:**
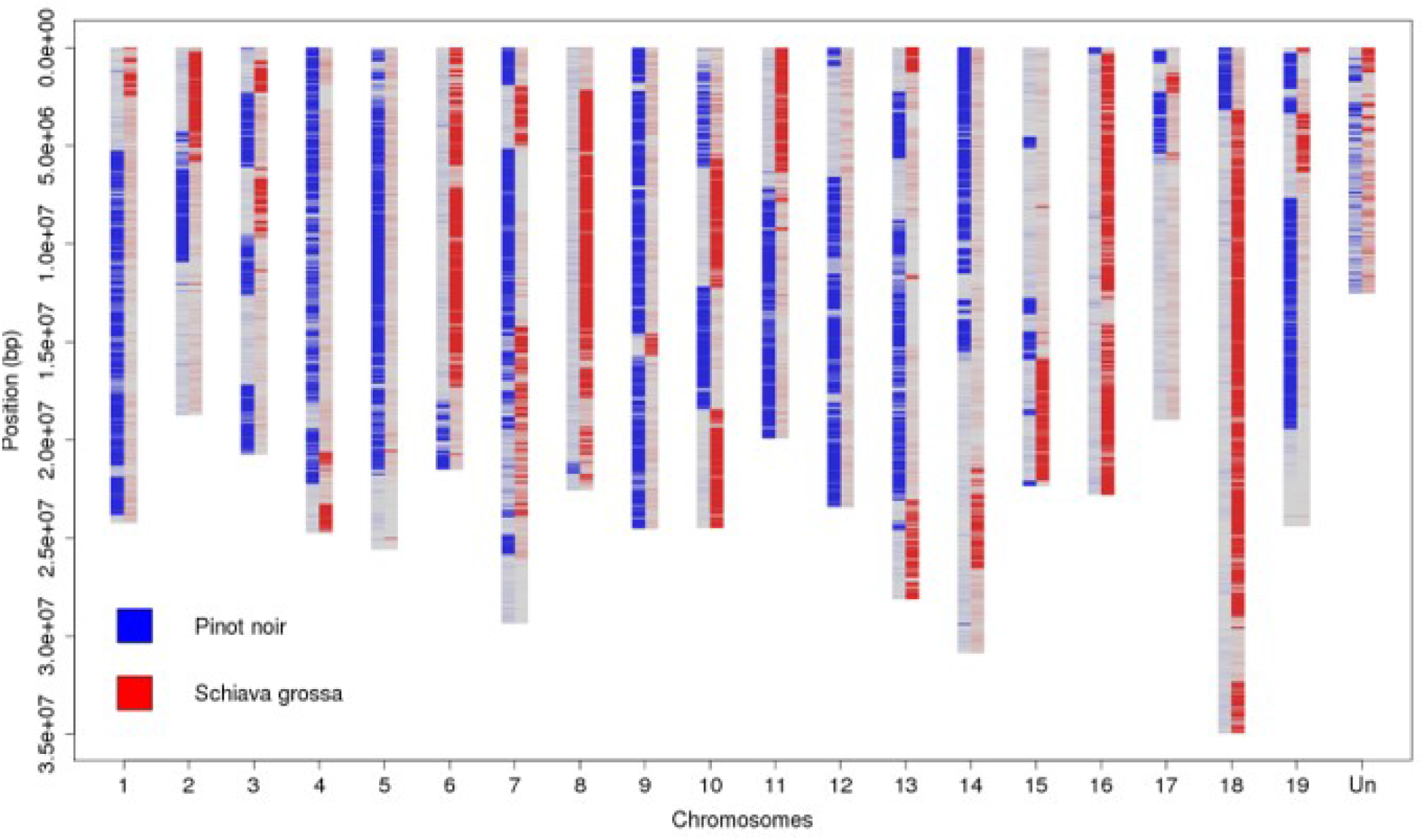
Density of ‘Pinot noir’ and ‘Schiava grossa’ homozygous SNPs compared to the PN40024.v4 genome assembly. The x-axis shows the 19 main pseudochromosomes and the artificial chrUn (‘Un’). The y-axis shows the base position in [bp]. Where density of ‘Pinot noir’ SNPs is high, it means PN40024.v4 carries the ‘Schiava grossa’ haplotype and *vice versa*. The regions where both ‘Pinot noir’ and ‘Schiava grossa’ SNP density is low correspond to regions where both genomes share a common haplotype.

**Fig. 5:**
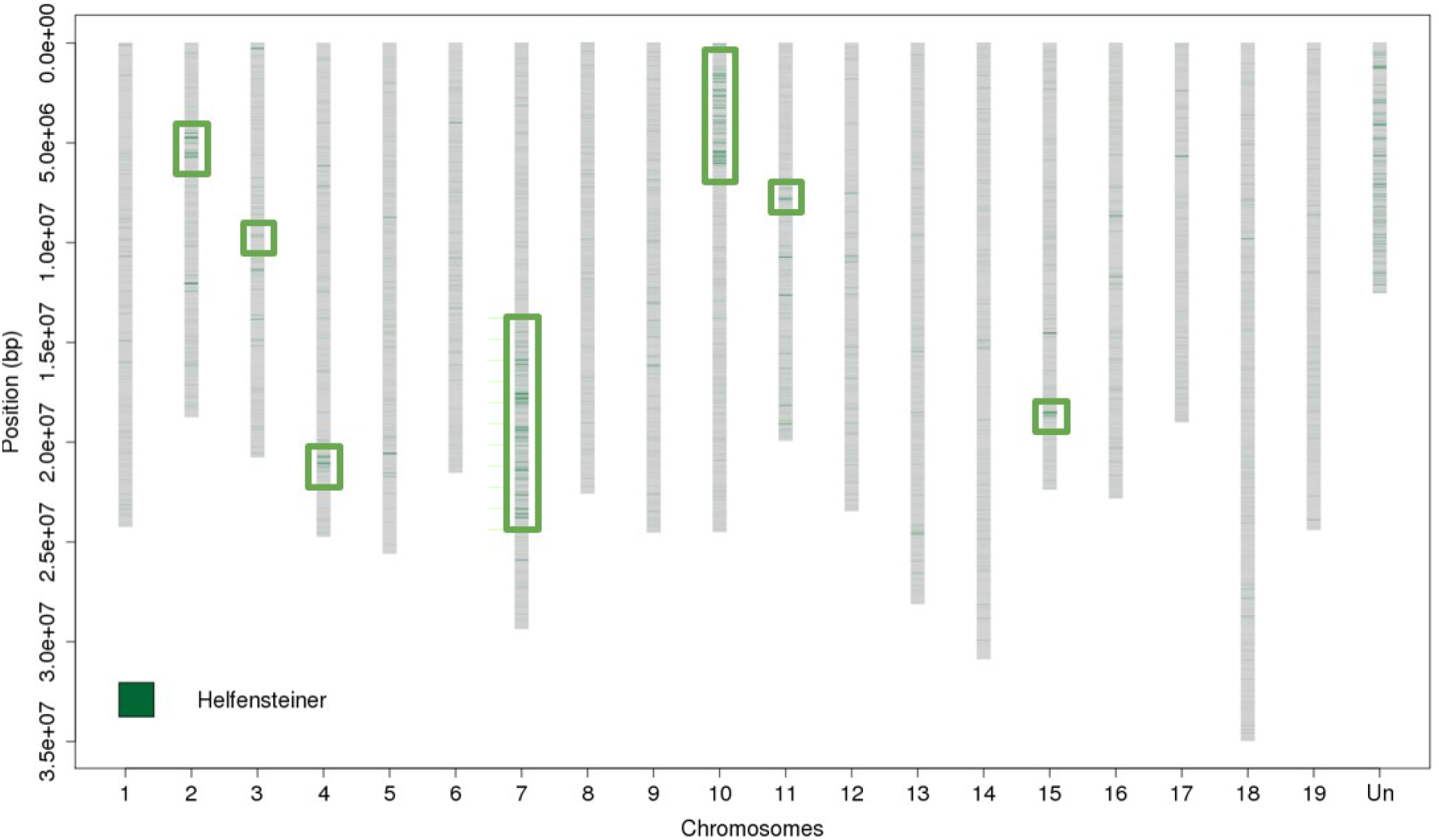
Density of ‘Helfensteiner’ homozygous SNPs compared to the PN40024.v4 genome assembly. The x-axis shows the 19 main pseudochromosomes and the artificial chrUn (‘Un’). The y-axis shows the base position in [bp]. The seven regions squared in green are the heterozygous regions.

**Fig. 6:**
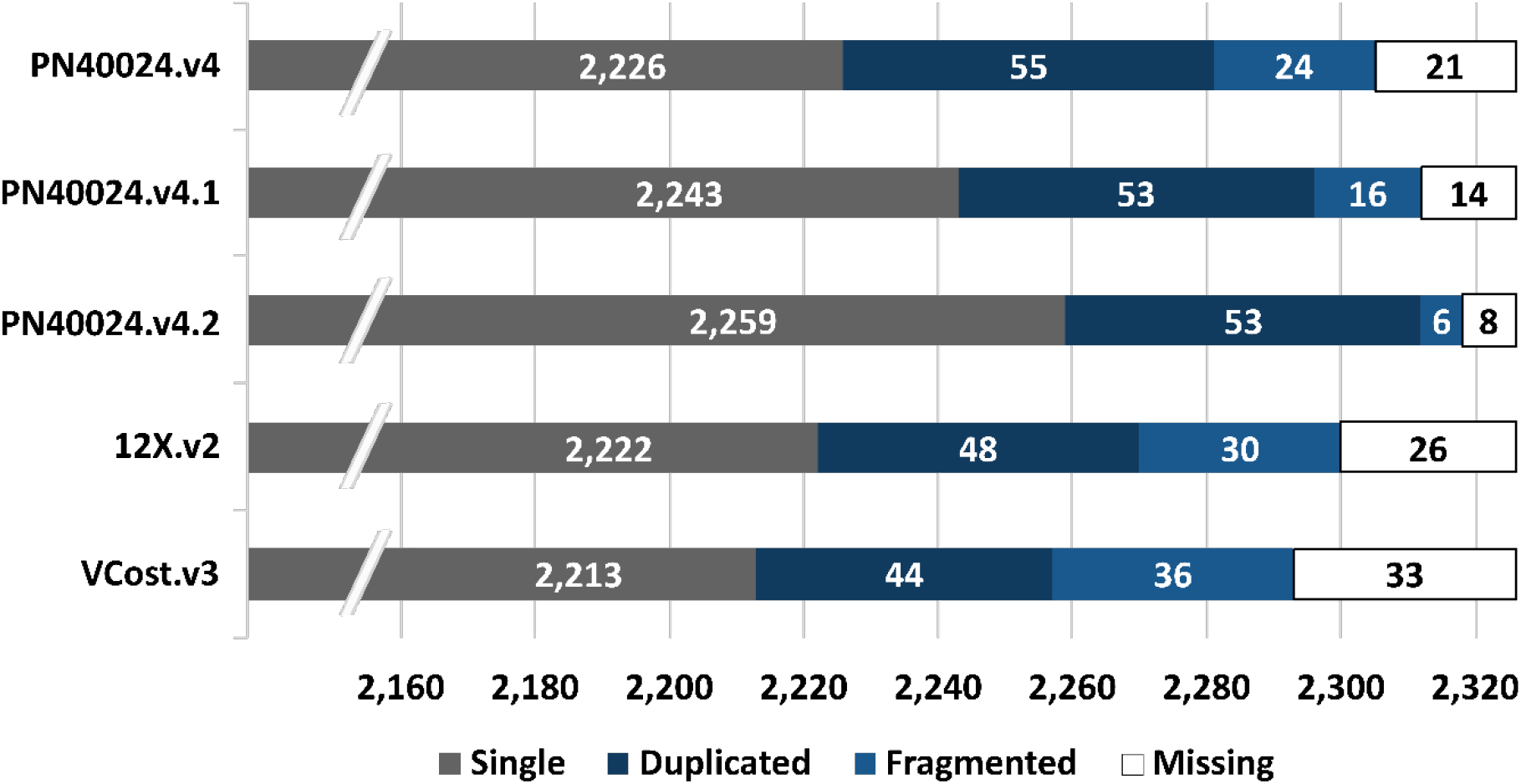
Plant core genes of the PN40024.v4 and PN12X.v2 genome assemblies and their annotations. The 2,326 plant core genes of the database eudicots_odb10 were determined in the PN40024.v4 genome assembly, in its annotation PN40024.v4.1, in the PN12X.v2 genome assembly and in the VCost.v3 gene annotation. ‘PN40024.v4.2’ is the PN40024.v4 gene annotation after manual curation of the fragmented and missing plant core genes.

The alignment metrics of PN40024 genomic Illumina paired-end reads were always better against PN40024.v4 compared to PN12X.v2, either for overall percentage of mapped reads (97.58% versus 96.58%) or for properly mapped pairs of reads (85.81% versus 82.82%) (Fig. 2). This confirms that the PN40024.v4 assembly is more complete and with a more accurate structure than PN12X.v2. Moreover, we compared alignments of 11 genomic Illumina paired-end read datasets from various cultivars against PN40024.v4 and PN12X.v2 assemblies, but also against ‘Cabernet Sauvignon’ ^5^ haplotype 1, whose assembly metrics and technology were similar to PN40024.v4. Again, PN40024.v4 performs best for each dataset, even when ‘Cabernet Sauvignon’ was aligned against its own assembly (Fig. 2). These results confirm that PN40024.v4 shows a quality suitable to become the new grapevine reference genome assembly, as it performs well with aligning genomic reads of various *V. vinifera* cultivars.

The error rate at nucleotide level was assessed by calling homozygous variations between PN40024 genomic Illumina paired-end reads aligned against the PN40024.v4 genome assembly. We identified 28.7 errors / Mb compared to 8.4 errors / Mb in the PN12X.v2 genome assembly. However, they are unevenly distributed along the chromosomes and they mostly co-localize with the newly assembled long read-based regions and the seven heterozygous regions (Fig. 3). A higher density of errors was also detected in the heterozygous regions of the PN12X.v2 genome assembly (Supplementary File 1 Fig. S2). We detected 284.4 errors / Mb in PN40024.v4 heterozygous regions of and 83.1 errors / Mb in PN12X.v2 heterozygous regions, which is, respectively, about 10 times denser than their average error rate. Thus, the overall increase of error rate in the PN40024.v4 assembly is mostly due to the use of SMRT long reads to improve the completeness of the reference genome assembly.

Using Merqury, the base level quality value (QV) of the PN40024.v4 genome assembly was estimated to be 36.02, which is slightly worse than QV of 37.43 of the PN12X.v2 genome assembly (Table 2). This result confirms that additional SMRT sequences are not as accurate as Sanger-based sequences and they slightly decrease overall accuracy of the assembly. Also, the error rate of the PN40024.v4 genome assembly was increased by 0,00006964% compared to PN12X.v2, but still represents an accuracy of 99.999749801%, a metric associated with high-quality genome assemblies.

**Table 2:** Assembly quality values of PN40024.v4 and 12X.v2. Assembly quality values measured by Merqury for PN40024.v4 and 12X.v2 genome assemblies. QV denotes base level quality value.

|  | **12X.v2** | **PN40024.v4** |
| --- | --- | --- |
| **QV** | 37.4338 | 36.0171 |
| **Error rate [%]** | 0.000180559 | 0.000250199 |
| **k-mer completeness [%]** | 96.79 | 96.96 |

Nevertheless, the k-mer completeness was raised from 96.79% to 96.96% for the PN40024.v4 assembly. Based on k-mer profiles of PN40024 and its parents (see “The origin of the PN40024 genotype” section for details), Merqury computed the inheritance spectrum (Supplementary File 1 Fig. S3) showing a low portion of read-only missing k-mers that are unique for the child read set (paired-end short reads of PN40024). The few missing sequences are probably due to sequencing errors, k-mers of novel variations or contamination from microbiome in PN40024 short reads, indicating an almost fully-complete PN40024.v4 genome sequence assembly. Also, as the spectrum shows a single 2-copy peak around 12x and that no 1-copy peak was observed at half the size, the k-mer analysis supports the assumptions of an almost homozygous grapevine genotype.

### The origin of the PN40024 genotype

So far, the PN40024 genotype was supposed to be originally derived from cv. ‘Pinot noir’ ^1^. However, we found 1,415,200 homozygous variants between ‘Pinot noir’ and PN40024.v4 (versus 17,696 homozygous variants of PN40024 against its own assembly), meaning that ‘Pinot noir’ haplotypes were completely missing at these locations. These homozygous ‘Pinot noir’ variants were unevenly distributed along the chromosomes and formed blocks (Fig. 4). We identified that the haplotypes of unknown origins could be assigned to ‘Schiava grossa’ (synonyms: ‘Trollinger’ and ‘Frankenthal’) as already suspected by Jaillon and colleagues ^1^. There were 953,735 homozygous variants found between cv. ‘Schiava grossa’ and PN40024.v4 and the formed haplotype blocks were highly complementary to ‘Pinot noir’ haplotype blocks (Fig. 4). As a negative control, the same analysis was performed with cv. ‘Araklinos’ and 2,273,888 homozygous variants were identified, evenly distributed along the chromosomes (Supplementary File 1 Fig. S4).

Using Merqury, only a small portion of hap-mer specific k-mers (parental specific k-mers of the assembled F1) were found in the PN40024.v4 genome assembly (Supplementary File 1 Fig. S3). With the use of read data from both parents and child, Merqury was able to compute haplotype blocks by using the parental specific k-mers as anchors. A total of 1,454 haplotype blocks were computed for PN40024.v4 sequences with additional 289 haplotype blocks for alternative heterozygous sequence regions and 2,575 haplotype blocks for the 12X.v2 genome assembly (Table 3). The N50 was measured to 2.05 Mb (REF), 0.25 Mb (ALT) and 1.76 Mb (PN12X.v2). Compared to the PN12X.v2 genome assembly, PN40024.v4 presented less haplotype blocks, but comprised almost all bases showing a higher N50 value, *i.e*. its haplotype blocks are more contiguous.

**Table 3:** Haplotype block statistics of PN40024.v4 and 12X.v2. Phasing accuracy estimation of Merqury for PN40024.v4 and 12X.v2 genome assembly. ALT denotes alternative heterozygous sequence parts of PN40024.v4.

|  | **12X.v2** | **PN40024.v4** | **ALT** |
| --- | --- | --- | --- |
| **Number of blocks** | 2,575 | 1,454 | 289 |
| **Total bases in blocks [bp]** | 474,845,411 | 468,703,133 | 19,519,697 |
| **Block N50 size [kb]** | 1,762 | 2,050 | 250 |
| **Switch error rate [%]** | 0.766002 | 0.959042 | 4.75944 |

A greater amount of paternal (‘Schiava grossa’) than maternal (‘Pinot Noir’) specific k-mers were identified. After identifying the origin of each haplotype block using segmentation, it is estimated that 41% of the genome harbours a ‘Schiava grossa’-specific haplotype and 27% a ‘Pinot noir’-specific haplotype. It is estimated that 32% of the genome shares a common haplotype between the two parents, *i.e*., that these regions could originate either from ‘Pinot noir’ or ‘Schiava grossa’ indicating that ~57% could originate from ‘Schiava grossa’ and ~43% from ‘Pinot noir’.

The switch error rate was determined to 0.96% (REF), to 4.76% (ALT) and to 0.77% (PN12X.v2). Some of the switches are probably due to sequencing errors in the additional long read-based sequences. Moreover, as the error rate of ALT sequences was measured to ~4.76%, portions of the alternative sequences are a mixture of the maternal and paternal haplotype, confirming that despite the improved separation of the two haplotypes in PN40024.v4, phasing is still not perfect.

By exploring the *VIVC* database (www.vivc.de), the ‘Helfensteiner’ cultivar was found to originate from a cross between ‘Pinot noir precoce’ (a clone of ‘Pinot noir’) and ‘Schiava grossa’. By performing the same variant calling analysis, 53,671 homozygous variants were found between cv. ‘Helfensteiner’ and PN40024.v4, with 543 homozygous variants / Mb in the heterozygous regions and 93 homozygous variants / Mb in the homozygous regions (Fig. 5). As a negative control, ‘Araklinos’ showed 3,967 homozygous variants / Mb in the heterozygous regions and 4,818 homozygous variants / Mb in the homozygous regions). Thus, the ‘Helfensteiner’ homozygous variants are almost six times denser in error-prone regions of the PN40024.v4 assembly, which makes them probable “false positive” homozygous variants. Apart from heterozygous regions, no blocks of homozygous variants could be identified, meaning that one of the two ‘Helfensteiner’ haplotypes is always present in the PN40024 genome. This confirms that the ‘Helfensteiner’ variety is the true parent of the first selfing, from which the PN40024 genotype was created after eight more selfings.

### PN40024.v4.1 gene prediction, functional annotation and manual curation

The PN40024.v4.1 gene annotation of REF haplotype comprises 35,922 gene models of which 35,197 are protein-coding and 725 encode for tRNAs (Table 4). In particular, 1,572 novel protein-coding genes were annotated in the newly assembled long read-based regions. For heterozygous regions, 1,855 and 1,809 protein-coding genes were predicted for REF and ALT haplotypes, respectively (Table 5). Most genes were predicted on the ~11 Mb heterozygous region on chromosome 7 with 830 on the reference sequence and 792 on the alternative sequence followed by the ~5 Mb region on chromosome 10 with 650 and 623 protein-coding genes.

**Table 4:**
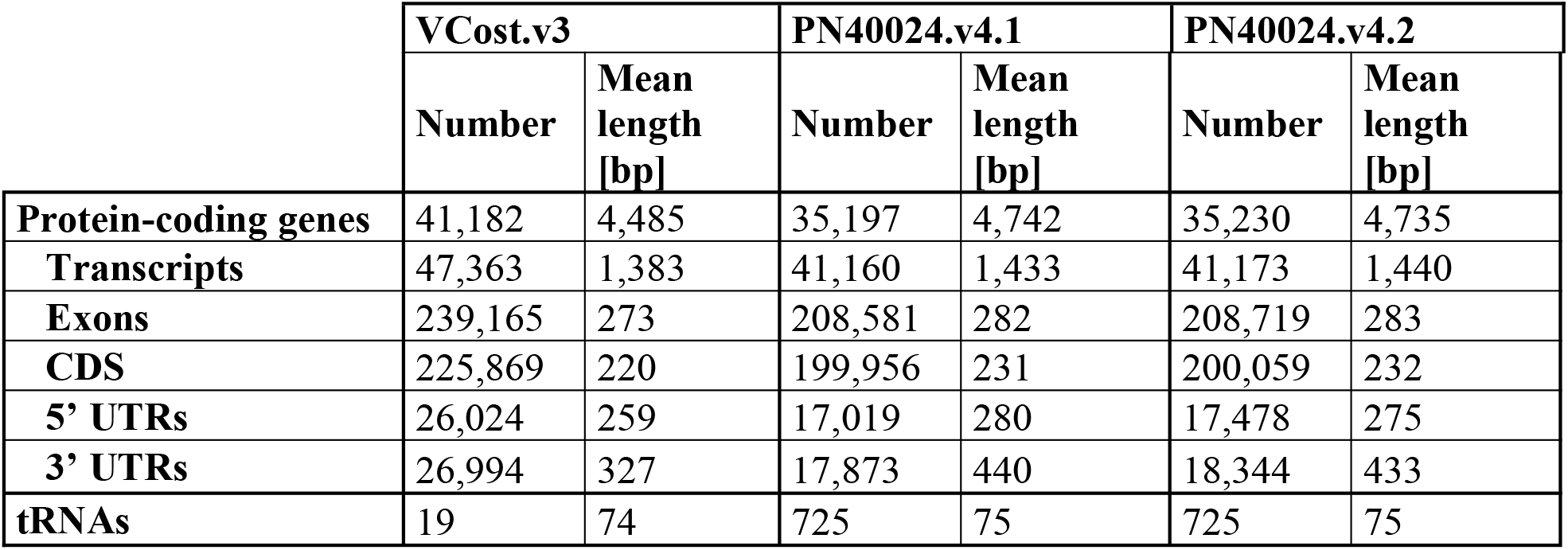
VCost.v3, PN40024.v4.1 REF haplotype and PN40024.v4.2 REF haplotype gene prediction overview.

**Table 5:**
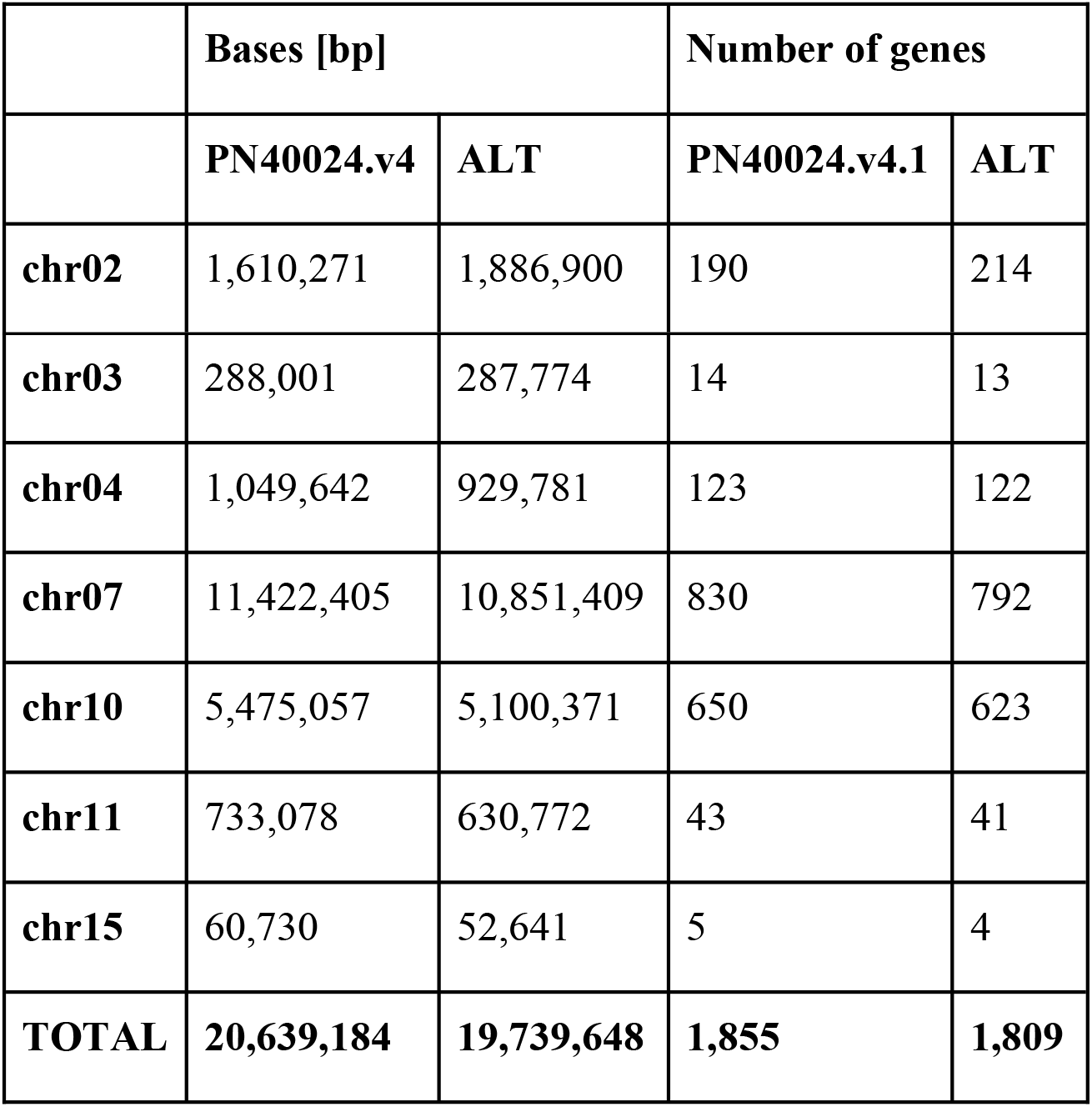
Gene numbers of heterozygous sequence regions. The abbreviation ALT denotes the alternative heterozygous sequence regions.

|  | Bases [bp] |  | Number of genes |  |
| --- | --- | --- | --- | --- |
|  | PN40024.v4 | ALT | PN40024.v4.1 | ALT |
| **chr02** | 1,610,271 | 1,886,900 | 190 | 214 |
| **chr03** | 288,001 | 287,774 | 14 | 13 |
| **chr04** | 1,049,642 | 929,781 | 123 | 122 |
| **chr07** | 11,422,405 | 10,851,409 | 830 | 792 |
| **chr10** | 5,475,057 | 5,100,371 | 650 | 623 |
| **chr11** | 733,078 | 630,772 | 43 | 41 |
| **chr15** | 60,730 | 52,641 | 5 | 4 |
| **TOTAL** | **20,639,184** | **19,739,648** | **1,855** | **1,809** |

To check for completeness of the gene models, the plant core genes of the database eudicots_odb10 were predicted with BUSCO (Fig. 6). Of the 2,326 searched plant core genes, 2,296 or 98.7% were classified as complete in the PN40024.v4.1 gene annotation. Only 16 were predicted as fragmented and only 14 were not found.

Compared to PN12X.v2 VCost.v3 gene annotation, PN40024.v4.1 counts less predictions (41,182 versus 35,197) but their size is longer on average (4,485 bp versus 4,742 bp) (Table 4). Also, the BUSCO analysis performed on VCost.v3 showed that 2,257 or 97.0% were classified as complete (Fig. 6). Thus, PN40024.v4.1 gene annotation represents PN40024 gene space in a more exhaustive and less fragmented manner compared to VCost.v3.

To help the community in the transfer of information across versions (*i.e*., correspondences), we retained as many gene names from VCost.v3 in PN40024.v4.1 as possible. We adopted a strategy based on RBHs followed by some filtering steps which allowed us to transfer names for 66% (23,206) of PN40024.v4.1 gene models with the nomenclature VitviXXg0YYYY (XX being the chromosome number and YYYY the number smaller than 4,000). One third (11,991) of PN40024.v4.1 gene models could not be named with a VCost.v3 identifier and were named with the nomenclature VitviXXg0**<u>4</u>**ZZZ (XX being the chromosome number and ZZZ the number smaller than 1,000). The detailed nomenclature for PN40024.v4.1 gene annotations is given in Supplementary File 2 Table S4.

The functional annotation of PN40024.v4.1 was performed using Blast2GO and resulted in at least one Gene Ontology term for 87% (30,689) of the genes and one Enzyme Code for 41% (14,512) of them. The main classes and ontologies are detailed in (Supplementary File 1 Fig. S5).

A subset of the RNA-Seq data published by ^81^ was used to compare the results of a differential gene expression analysis performed with PN12X.v2/VCost.v3 and PN40024.v4/PN40024.v4.1. In terms of mapping, the percentage of aligned reads was equivalent or slightly better when using PN40024.v4 genome assembly compared to PN12X.v2 (Supplementary File 2 Table S6). Additionally, the percentage of assigned reads, *i.e*., the percentage of reads aligned under an annotated gene, was 2.4 to 3% better with PN40024.v4/PN40024.v4.1 compared to PN12X.v2/VCost.v3, which confirms the improved quality of PN40024.v4.1 gene annotation. Moreover, after differential gene expression analysis, the use of PN40024.v4/PN40024.v4.1 allowed identifying more differentially expressed genes than PN12X.v2/VCost.v3 (Supplementary File 1 Fig. S6). This result along with the exhaustive functional annotation of PN40024.v4.1 shows that this new version of the PN40024 reference genome and annotation is a very efficient resource to perform transcriptomics and functional enrichment analyses.

Despite marked improvement of the PN40024.v4.1 automated annotation with respect to the previous VCost.v3 annotation, some recently expanded gene families have not been comprehensively annotated, such as the stilbene synthase (STS) gene family. Therefore, 1,641 genes (1,579 edited and 62 deleted) were manually curated using a purpose-built Apollo server (http://138.102.159.70:8080/apollo) providing a wide range of transcriptomic and genomic data for PN40024.v4. In an effort to preserve previous VCost.v3 manual curation and functional annotation efforts, a particular focus was given to genes present in the reference catalogue (Navarro-Payá et al, 2022). The PN40024.v4.1 automated annotation including the manually curated features was called PN40024.v4.2, which metrics are presented in (Table 4). An automated annotation from PN40024.v4.1 that was manually curated was deleted and replaced by its curated version in PN40024.v4.2. Also, same rules were applied for gene name transfer and nomenclature for PN40024.v4.1 and PN40024.v4.2. The BUSCO analysis performed on PN40024.v4.2 shows that the fragmented plant core genes were reduced to six and the missing genes to eight (Fig. 6). Thus, PN40024.v4.2 gene models comprise 2,308 or 99.2% complete plant core genes.

## Conclusion

The here provided PN40024.v4 assembly is the most suitable grapevine reference genome sequence assembly as it notably outperforms PN12X.v2. In terms of genomic and transcriptomic read mapping, the assembly also outperforms other high-quality *V. vinifera* genome assemblies, something that even occurs when reads from these recently sequenced cultivars are used. Having a fully resolved alternative haplotype sequence, more continuous sequences and resolving many up-to-now unknown bases, PN40024.v4 represents the almost complete diploid genome of the PN40024 genotype. Despite the many improvements and advances that PN40024.v4 has experienced, the genome sequence is still not perfect in regard to haplotype switching and to newly introduced errors by the implementation of long genomic reads. Further improvements should focus on these regions. Nevertheless, the gene annotation of PN40024.v4 should be used as the most updated resource for transcriptomics and functional enrichment analyses, while the genes of heterozygous regions that are likely represented on both haplotypes will allow exploring heterozygous genetic traits.

## Data availability and tools

Raw sequencing data and the PN40024.v4 genome assembly are available at ENA under BioProject PRJEB45423. Also, the PN40024.v4 genome assembly with structural and functional gene annotation is available on the INTEGRAPE website (https://integrape.eu/resources/genes-genomes/genome-accessions), on the Grape Genomics Encyclopedia portal (http://grapedia.org/) and under the DOI number doi:10.57745/F9N2FZ (https://entrepot.recherche.data.gouv.fr/dataset.xhtml?persistentId=doi:10.57745/F9N2FZ). A Sequence Server v2.0.0 interface (http://138.102.159.70:4567/) was set up to perform BLAST analyses. A JBrowse interface (http://138.102.159.70/jbrowse/) was set up to visualize PN40024.v4 assembly and PN40024.v4.1 and v4.2 annotations, but also some previous annotation versions that were transferred, some RNA-Seq alignments and miscellaneous tracks. An Apollo interface (http://138.102.159.70:8080/apollo; training and account mandatory) was set up to manually curate gene annotations according to the dedicated guidelines (https://integrape.eu/resources/data-management/).

## Acknowledgements

We thank INRAE department “Biologie et Amélioration des Plantes” for funding; experimental unit UEAV (INRAE, Colmar) for plant maintenance; Anne-Marie Digby (University of Verona) for English correction; Emilce Prado for plant DNA extractions; EPGV (INRAE, Evry) for library prep and DNA sequencing; CNRGV (INRAE, Toulouse) for long read DNA extractions; Gentyane platform (INRAE, Clermont-Ferrand) for SMRT sequencing; Dr. Timothée Flutre and Amandine Launay for their help in coordinating the FruitSelGen project and in acquiring GBS data; Mario Pezzotti, Anne-Françoise Adam-Blondon, Michele Morgante, Gabriele Di Gaspero and Gabriele Magris for helpful discussions during the project; Pablo Carbonell-Bejerano (Instituto de Ciencias de la Vid y el Vino - ICVV) for critically reviewing the manuscript. This work was supported by the BMBF-funded de.NBI Cloud within the German Network for Bioinformatics Infrastructure (de.NBI). This article is based upon work from COST Action CA17111 INTEGRAPE, supported by COST (European Cooperation in Science and Technology).

## Supplementary information

Supplementary File 1 Fig. S1: Overview of PN40024 assembly and annotation versions Supplementary File 1 Fig. S2: Density of errors and of heterozygous SNPs in the PN12X.v2 assembly

Supplementary File 1 Fig. S3: Inheritance spectrum plot of PN40024.v4

Supplementary File 1 Fig. S4: ‘Araklinos’ homozygous SNP density in the PN40024.v4 genome assembly

Supplementary File 1 Fig. S5: Gene Ontology results

Supplementary File 1 Fig. S6: Gene Ontology results level 3

Supplementary File 1 Fig. S7: Comparison of results of the differential gene expression analysis between PN12X.v2 and PN0024.v4

Supplementary File 2 Table S1: Stranded *V. vinifera* RNA-seq data

Supplementary File 2 Table S2: Unstranded *V. vinifera* RNA-seq data

Supplementary File 2 Table S3: Datasets and weights used for data combination with EvidenceModeler

Supplementary File 2 Table S4: Gene identifier nomenclature and transfer

Supplementary File 2 Table S5: Assembly and anchoring statistics for the second allelic version, ALT, of PN40024.v4

Supplementary File 2 Table S6: Comparison of RNA-Seq read alignments to the PN12X.v2 and PN40024.v4 assemblies and assignment to VCost.v3 and PN40024.v4.1 gene predictions

