## Supplementary File 1 Figures for "An improved reference of the grapevine genome supports reasserting the origin of the PN40024 highly-homozygous genotype"

### Supplementary Figures

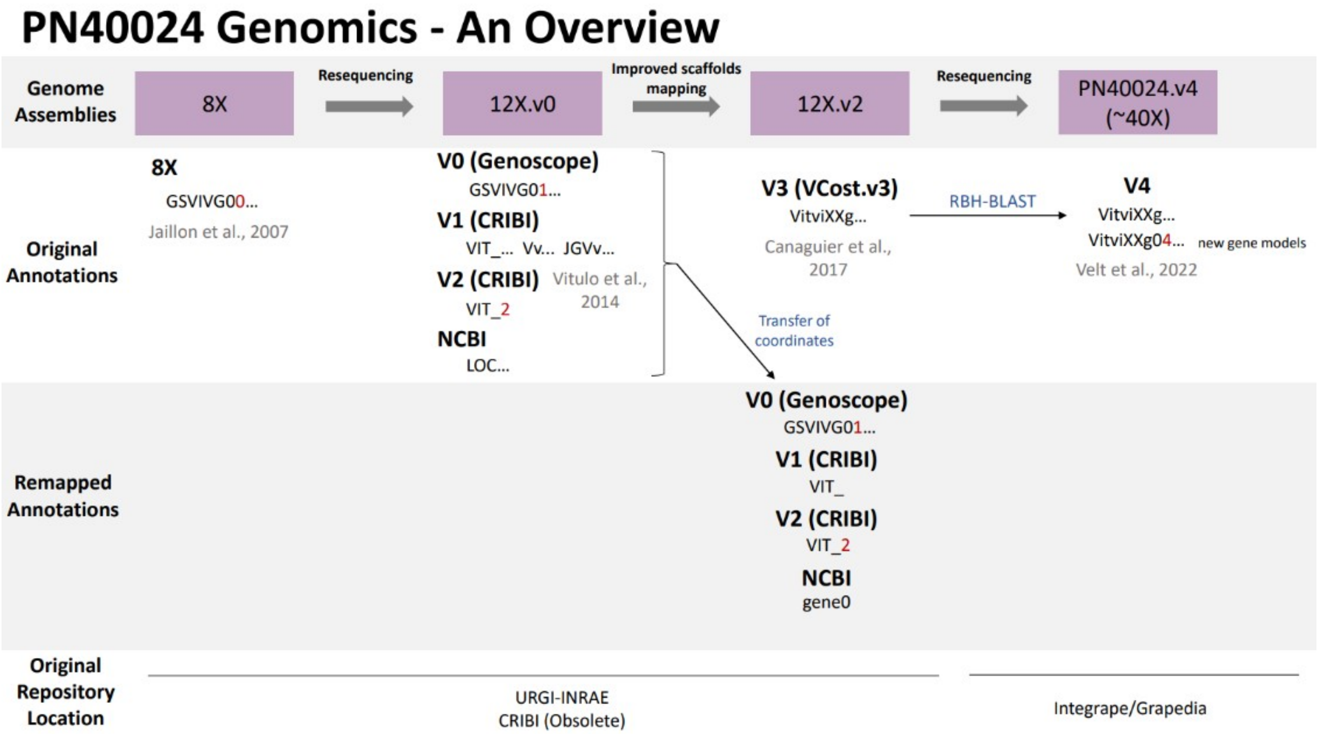


**Fig. S1 – Overview of PN40024 assembly and annotation versions.**

The PN40024 genome has been improved since its release in 2007 and has resulted in four different genome assemblies so far. Each genome version has at least one original gene annotation version associated to it. Moreover, some annotation versions have been transferred to newer genome assemblies. Gene ID prefixes for each annotation are indicated with key differences shown in red.

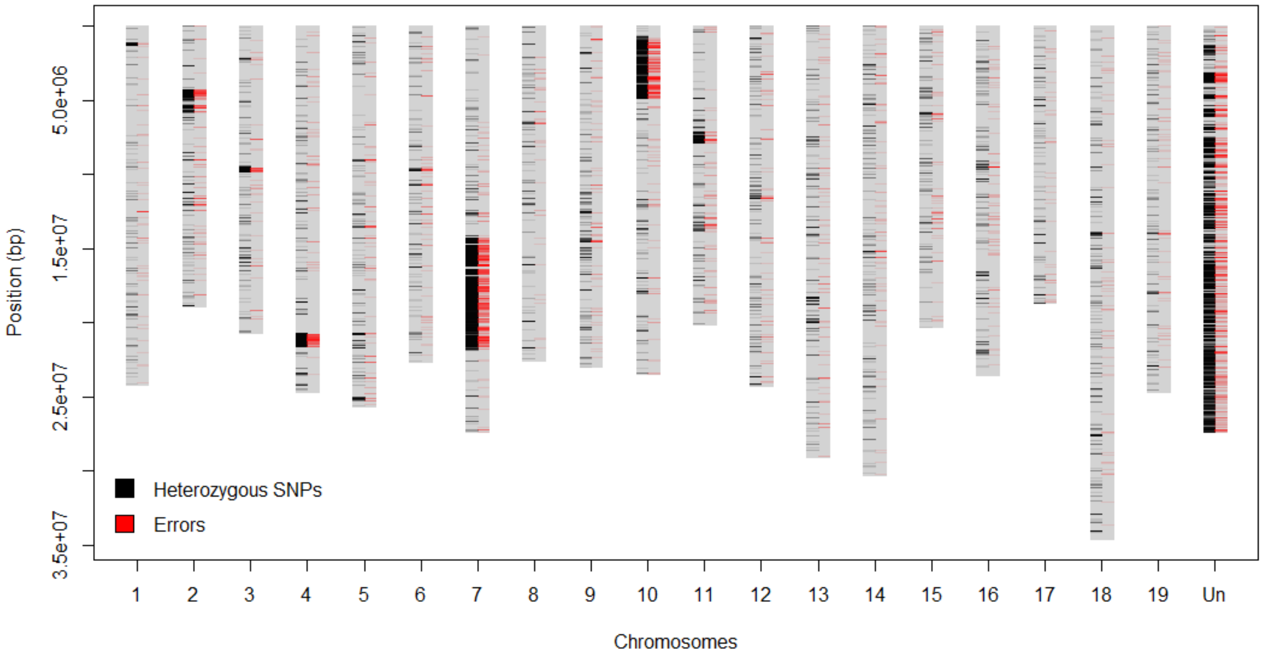


**Fig. S2 – Density of errors and of heterozygous SNPs in the PN12X.v2 assembly.**

The x-axis shows the 19 main pseudochromosomes and the artificial chrUn ('Un'). The y-axis shows the base position in [bp].

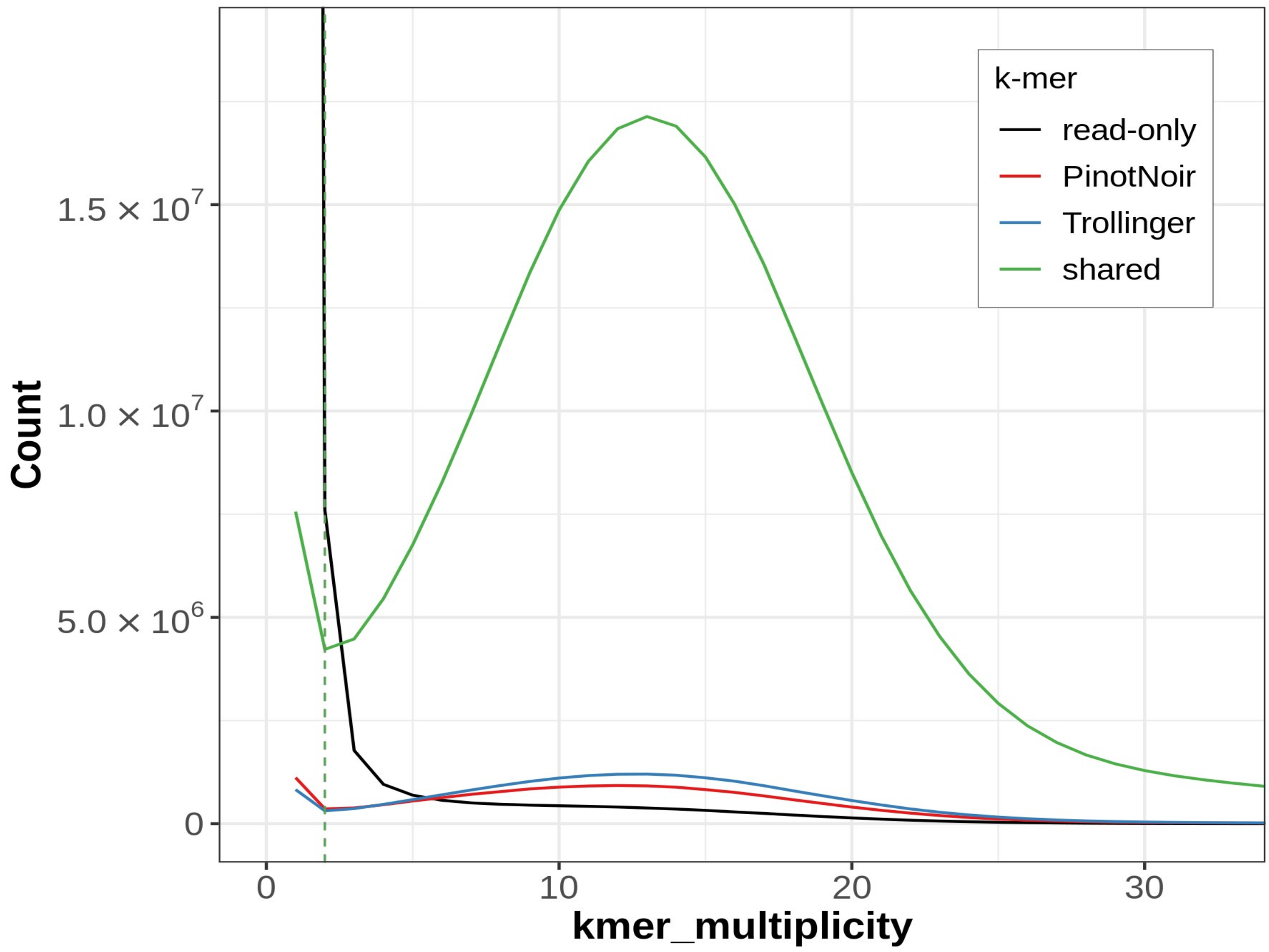


**Fig. S3 – Inheritance spectrum plot of PN40024.v4.**

The plot shows the k-mer multiplicity of the PN40024 (child) read set separated by inheritance. k-mers uniquely found in the child read set are colored black, k-mers specific for the paternal ‘Schiava Grossa’ (‘Trollinger’) read set are colored blue, k-mers specific for the maternal ‘Pinot Noir’ read set are colored red and k-mers found in the read sets of both parents are colored green.

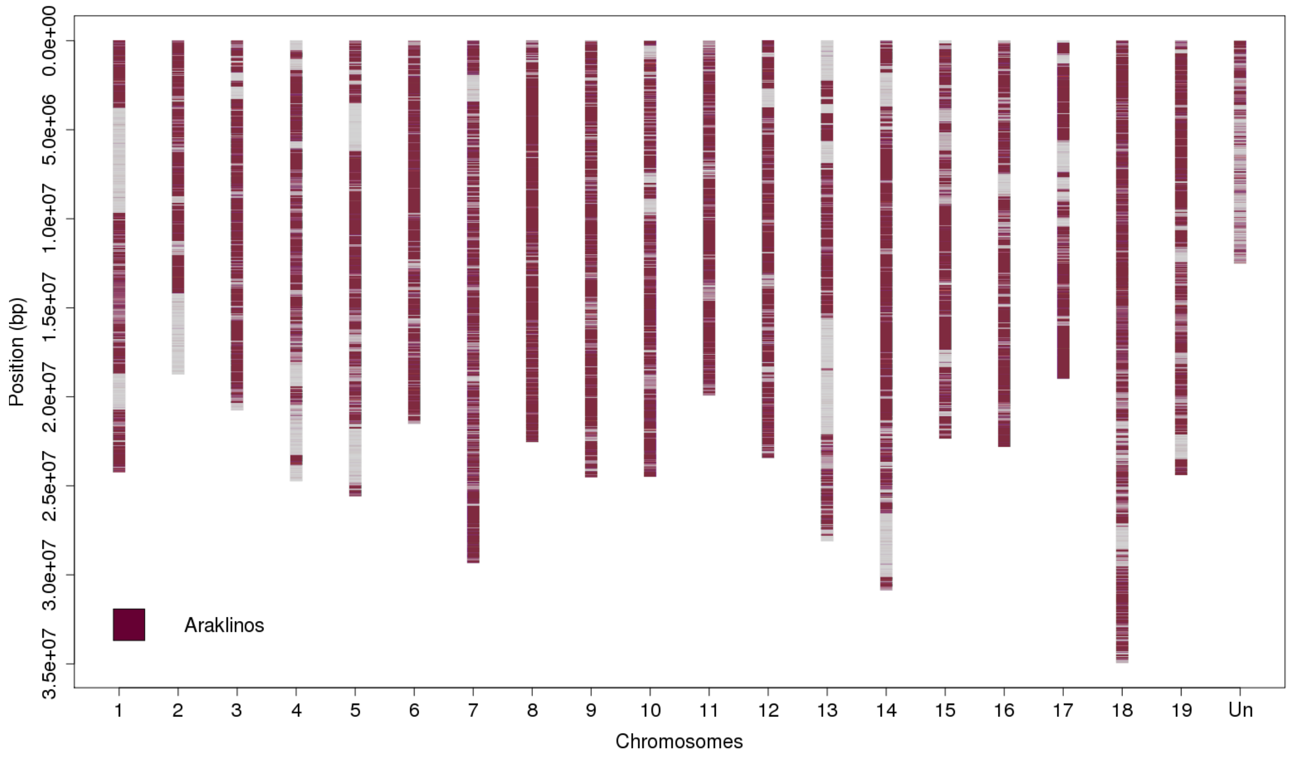


**Fig. S4 – 'Araklinos' homozygous SNP density in the PN40024.v4 genome assembly.**

The x-axis shows the 19 main pseudochromosomes and the artificial chrUn ('Un'). The y-axis shows the base position in [bp].

A

B

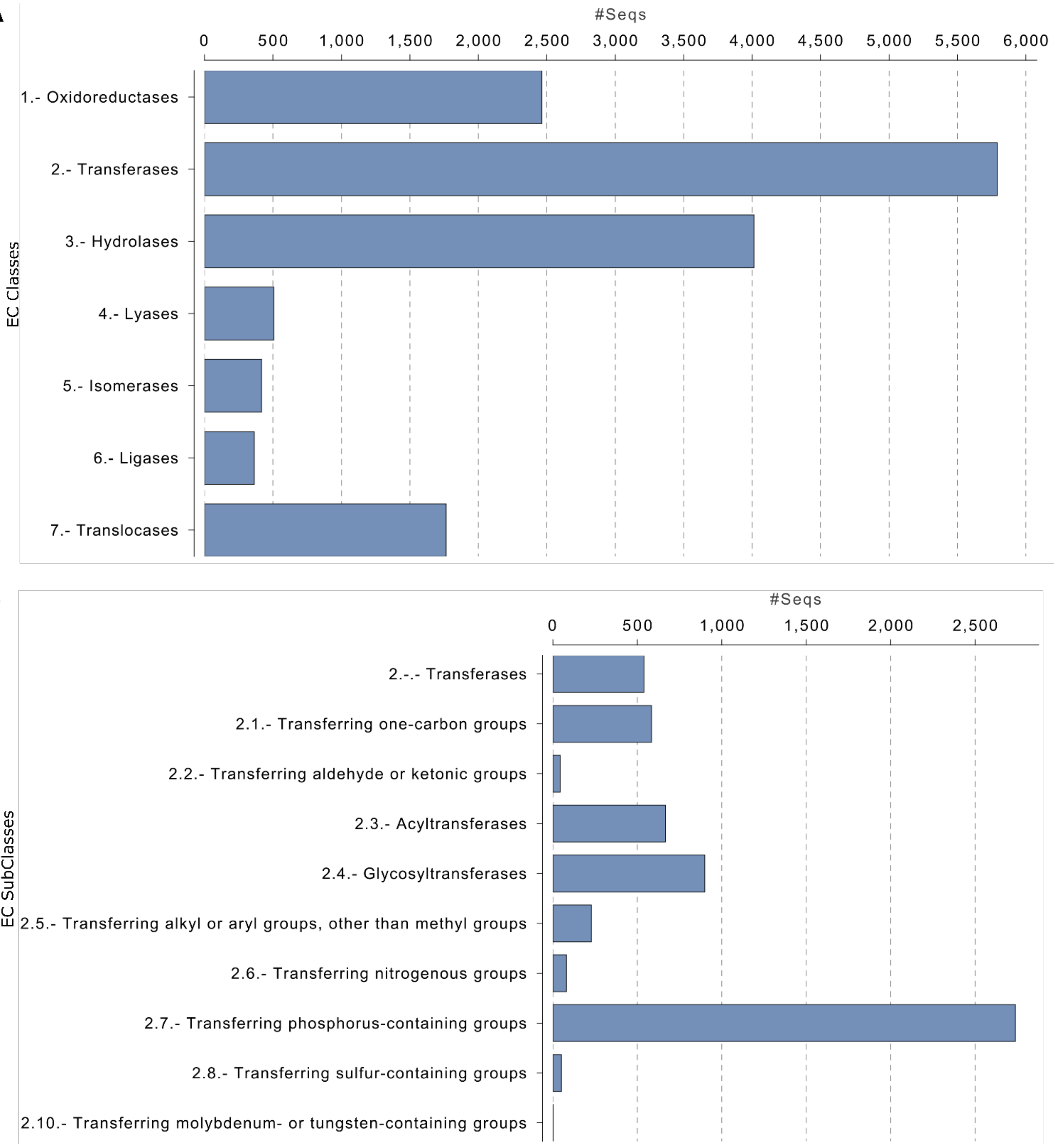


**Fig. S5 – Gene Ontology results.**

Plots are part of Blast2GO report. **A**) Main enzyme code level distribution. **B**) Enzyme code level 2 distribution.

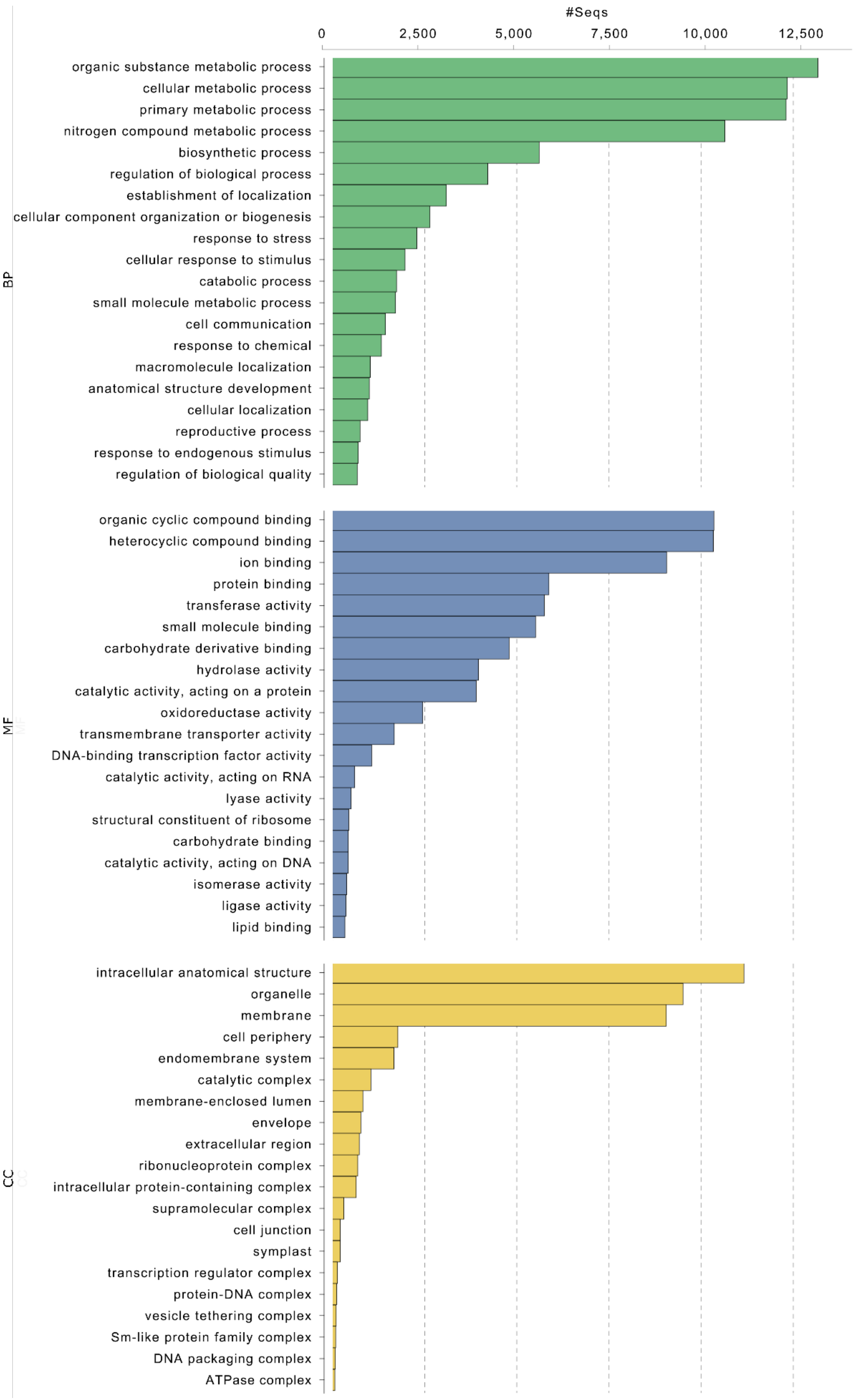


**Fig. S6 – Gene Ontology results level 3.**

GO distribution in level 3 (Top 50). Plot is part of Blast2GO report.

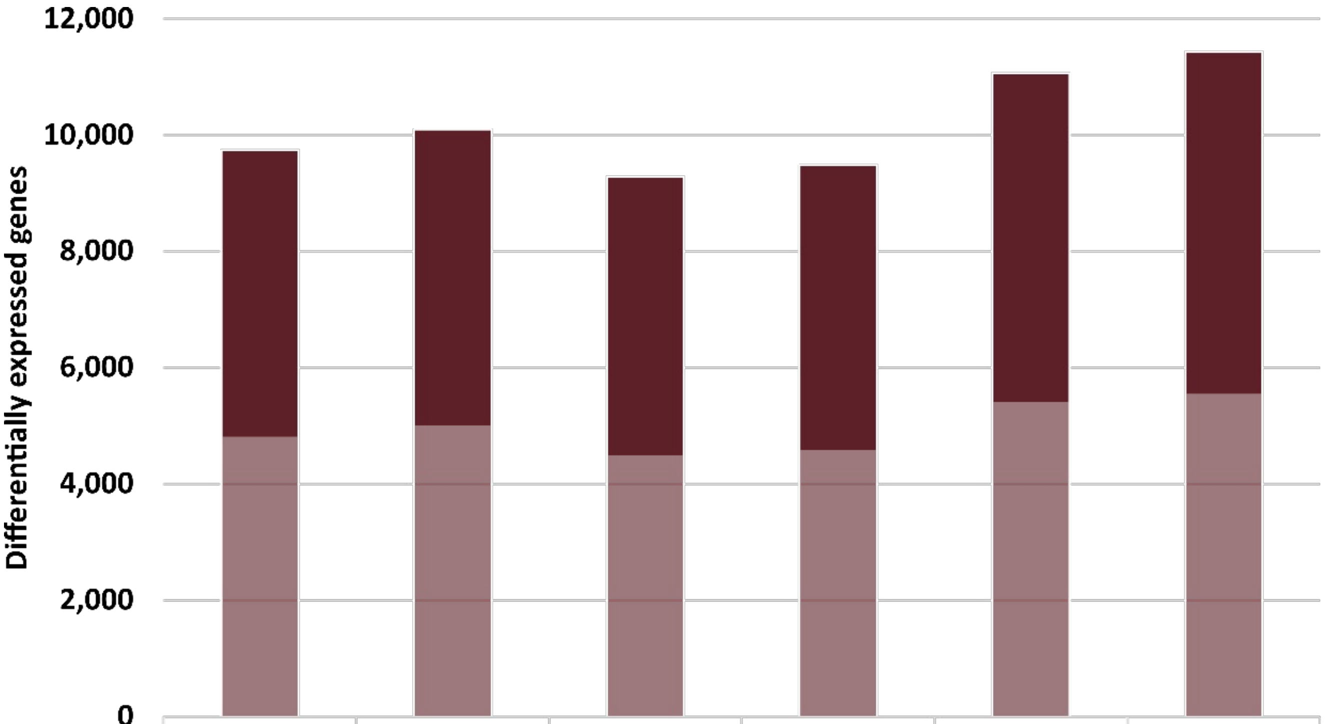


| | 'Barbera' vs. 'Sangiovese' PN12X.v2 | 'Barbera' vs. 'Sangiovese' PN40024.v4 | 'Refosco' vs. 'Sangiovese' PN12X.v2 | 'Refosco' vs. 'Sangiovese' PN40024.v4 | 'Refosco' vs. 'Barbera' PN12X.v2 | 'Refosco' vs. 'Barbera' PN40024.v4 |
|---|---|---|---|---|---|---|
| Up | 4,930 | 5,089 | 4,786 | 4,900 | 5,669 | 5,890 |
| Down | 4,808 | 5,008 | 4,493 | 4,589 | 5,403 | 5,548 |
| Sum | 9,738 | 10,097 | 9,279 | 9,489 | 11,072 | 11,438 |

**Fig. S7 – Comparison of results of the differential gene expression analysis between PN12X.v2 and PN0024.v4.**

Shown are the number of differentially expressed genes (DEGs) determined for the PN12X.v2 genome assembly with VCost.v3 gene annotation or for the PN40024.v4 assembly with PN40024.v4.1 annotation. 'Barbera', 'Sangiovese' and 'Refosco' denote the used RNA-Seq datasets. 'Up' refers to up-regulated DEGs (dark red) and 'Down' to down-regulated DEGs (pale red).
